# Continuous evolution of gene libraries towards arbitrary functions reveals the versatility of biomolecular evolution *in vivo*

**DOI:** 10.1101/2025.03.22.644768

**Authors:** Alexander (Olek) Pisera, Alireza Tanoori, Chang C. Liu

## Abstract

Life relies on constant biomolecular innovation, but the slowness of natural evolution limits our ability to witness new gene functions emerging in real time. Consequently, much remains unknown about how – and how easily – functional biomolecules originate and evolve. To address this gap, we encoded diverse gene libraries (*e.g.*, open reading frames from *Saccharomyces cerevisiae* and *Escherichia coli*) onto an orthogonal DNA replication (OrthoRep) system and exposed the resulting continuously hypermutating genetic repertoires to a multitude of complex selection pressures *in vivo*. This experimental template mimics the “multi-gene, multi-objective” possibility landscape of natural evolution, but at drastically accelerated evolutionary speed. In only ∼100 generations (∼1 month), populations evolved an abundance of novel gene functions when challenged with diverse pressures including vitamin deficiency, metal toxicity, promoter inactivity, and protein degradation. Notably, several evolved outcomes originated from sequences with no initial effect on fitness, demonstrating both emergence and evolution of biomolecular function. Evolved outcomes proved highly potent, often increasing fitness by several-fold under selection. Evolved genes spanned a range of molecular functions including *de novo* metal binding, transcription factor activity, and modulation of protein degradation, as supported by biochemical and RNA-seq experiments. Overall, our findings show that novel gene functions can originate with surprising ease in cellular contexts and motivate further continuous evolution experiments that agnostically explore the fertility of sequence space *in vivo*.

## Introduction

How new gene functions emerge and evolve is a fundamental question in biology. Dominant models include neofunctionalization, which poses duplication of existing genes and divergence as the key mechanism (*1–6*), as well as *de novo* gene birth, which posits that non-coding sequences, intronic sequences, or alternative frames of existing ORFs become (aberrantly) expressed and enter the view of selection (*7–14*). However, experimentally testing these mechanisms and probing the broader origins questions of biomolecular function is challenging because evolution is slow. For example, in a landmark experimental demonstration of a neofunctionalization process, it took 3000 generations for a histidine biosynthesis enzyme (HisA) to evolve weak tryptophan biosynthesis (TrpF) activity, duplicate, and diverge into a gene with strong TrpF activity (*15*). Furthermore, HisA and TrpF share a high degree of relatedness, carrying out the same isomerization reaction (on different substrates) and naturally descending from a generalist ancestor, so the biomolecular innovation reached here was arguably modest and chosen based on its expected feasibility (*16*). To explore whether existing genes can begin evolving towards *arbitrary* functions in shorter experiments, synthetic libraries of *Escherichia coli* genes have been subjected to selections for antibiotic and toxin resistance, producing a large number of examples where an overexpressed gene was able to directly or indirectly confer some degree of resistance (*17*). Likewise, experiments that tested the evolutionary productivity of classically “non-coding” or *de novo* generated genes of varying degrees of randomness (*18–28*) support the idea that perhaps more genetic matter than previously thought can be the origin of new genes. However, few of these efforts also demonstrate the evolutionary improvement of newly discovered genes (*22*, *23*, *26*), leaving the fitness effects of new genes weak and accounts of their evolutionary potential incomplete. Therefore, our understanding of how – and how easily – biomolecular functions come to be remains fragmented.

Nevertheless, a compelling picture is emerging where 1) the wealth of genetic material that can evolve after duplication, horizontal gene transfer, or gene birth, 2) the commonness of secondary activities in extant genes (*e.g.*, moonlighting (*29–31*) or promiscuous (*32–35*) activity), and 3) the systems level complexity of cells that can afford many ways to achieve selectable phenotypes, may combine to make the evolution of novel gene functions far less constrained than commonly thought. Recognizing this confluence, we present OrthoRep (*36*, *37*) Assisted Continuous Library Evolution (ORACLE) (**Fig. 1**) as a general strategy to prospectively study how large genetic repertoires can originate and evolve novel gene functions *in vivo*. In ORACLE, highly diverse, multi-length libraries of genes (*e.g.*, comprehensive collections of *E. coli* and *Saccharomyces cerevisiae* protein-coding sequences) are encoded onto the p1 plasmid of OrthoRep, which accelerates their continuous mutation by up to 1-million-fold over genomically encoded genes in *S. cerevisiae* (*38–40*). The resulting populations are then propagated under a range of selection pressures, mimicking the fact that in natural evolution, any number of genes can be the source of innovations that satisfy any number of selection pressures. In evolution experiments lasting only ∼100 generations (∼1 month), ORACLE yielded many examples where new genes evolved to satisfy the imposed selection pressures. In the most striking cases, genes without known functions (and whose wild type (WT) sequences had no effect on fitness) evolved to satisfy selection, including examples where a frameshift-generated peptide sequence originated a novel metal-binding function and where cryptic prophage genes evolved transcription factor and protein homeostasis modulation activities *de novo*. By subjecting diverse, function agnostic, variable length, high complexity genetic libraries to continuous directed evolution *in vivo*, ORACLE allows us to witness the flexibility of *in vivo* gene evolution in real time.

**Fig. 1.**
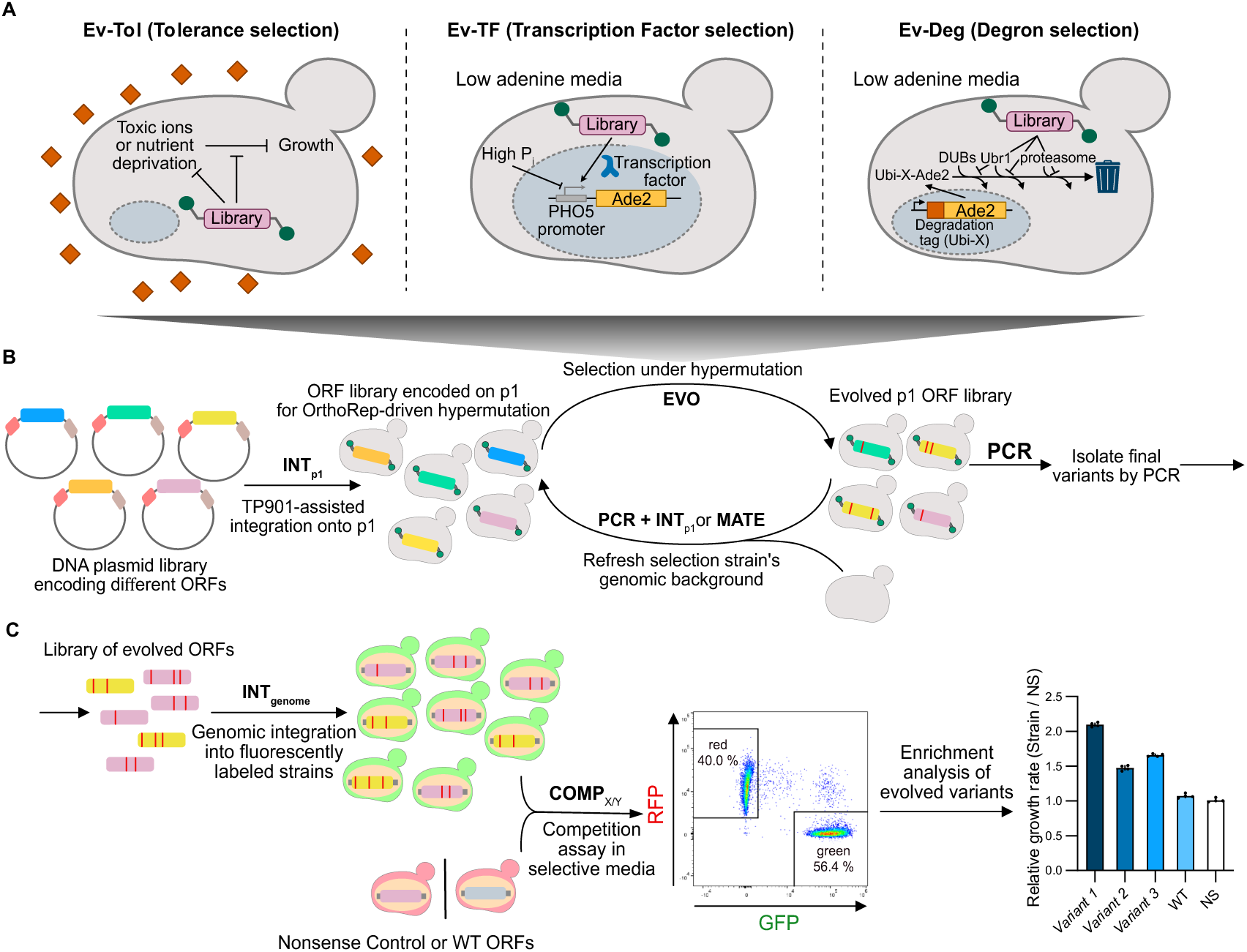
ORACLE expands the capabilities of OrthoRep to library evolution. **(A)** Overview of main selection schemes used for evolution experiments in this study. In ***Ev-Tol***, we utilized non-optimal growth media to evolve genes that overcome vitamin deprivation or metal toxicity. ***Ev-TF*** links cell survival to transcriptional regulation, specifically selecting for activation of a repressed phosphate-starvation-response promoter controlling the expression of the *ADE2* gene that is essential in media lacking adenine. In ***EV-Deg***, we employed an N-end-destabilized *ADE2* selection marker to select for the disruption of Ubr1-mediated degradation of N-end substrates or other protein degradation processes. **(B)** Schematic of ORACLE evolution. In ORACLE, gene libraries are first transformed onto p1 with **INT**_p1_ and subjected to an arbitrary evolutionary pressure under hypermutation. Gene libraries are evolved with frequent refreshing of genome, utilizing either **MATE** or **PCR + INT**_p1_. After several rounds of evolution and refreshing the genome, the genes are taken to the **COMP** stage for assessment. **(C)** In **COMP**, the genes evolved by ORACLE are assessed for their fitness advantage in a pooled outgrowth experiment where the ratio of cells containing evolved genes and control constructs are monitored over time by flow cytometry during growth in selection conditions. Typically, the library of evolved genes is integrated into a fluorescently labeled strain (**INT**_genome_) so that each cell encodes a single copy of an evolved gene. Then, the mixed population is grown in competition against reference cells, encoding a different fluorophore, with a nonsense (NS) gene is integrated. If populations outcompete reference NS cells, individual evolved gene variants are isolated and grown in competition against the NS cells and further competed against the wild type variant of the evolved gene (WT). In all competition experiments, the ratio of growth rates of one fluorescent strain against the other is obtained by tracking the expansion of the total population and the change in the ratio of the two strain types, as counted by flow cytometry. The exact values of the resulting bars represent the ratio of Malthusian growth rates between the two strain types, assuming exponential growth.

## Results

### Overview of an ORACLE experiment

In this work, we carried out 55 ORACLE experiments (**Table S1**), where an experiment is defined as a distinct gene library from which we attempted to evolve a particular function (see next section). Each experiment was carried out in at least two variations, one with the error-prone DNAP Trixy (error rate of 4 x 10^-5^ substitutions per base (s.p.b.)) and the other with the error-prone DNAP BadBoy3 (error rate of 10^-4^ s.p.b.) driving hypermutation of the p1-encoded gene library (*40*). Evolution experiments started from three gene libraries: 1) a pooled yeast ORF library (*41*) wherein *ADE2*, *TRP5*, *URA3*, *HIS3*, *KAR1*, *LYS2*, *CAN1*, *LEU2*, *LYP1*, *MET15* were omitted due to our reliance on these genes as auxotrophic markers and selection systems, 2) a comprehensive *E. coli* ORF library where each ORF’s C-terminus was fused to GFP (*42*, *43*), and 3) the same *E. coli* ORF library without the GFP fusion, which we generated by subcloning. (See **Supplementary Text** and **Materials and Methods** for additional information on library characteristics and construction.)

Each evolution experiment began with the “subroutines” **PCR**_lib_ and **INT**_p1_ to install a gene library into a strain carrying the appropriate genomic background for the intended functional selection. (See **Supplementary Text** and **Fig. S1-S6** for detailed descriptions of experimental subroutines, such as **PCR**_lib_, **PCR**, **INT**_p1_, **INT**_genome_, **MATE**, **COMP**_pop/NS_, **COMP**_pop/WT_, **COMP**_mut/NS_, and **COMP**_mut/WT_, how they were developed and optimized for the specific demands of ORACLE, and their experimental validation. We reference these subroutines with the designated shorthand throughout.) Each strain has either of two error-prone DNAPs, Trixy and BadBoy3 (*40*), encoded to drive the gene library’s hypermutation. Each population was then serially passaged in the corresponding selective condition with adjustment of the population size transferred at each passage based on optical density (OD) measurements. This OD-based monitoring allowed us to coordinate dozens of evolution experiments (**EVO**) in parallel on the same schedule and was also used to guide how we modified the selection strength. After about 4 passages under selection, evolving p1 populations were transferred into fresh strains using **MATE** or **PCR** + **INT**_p1_ (**Supplementary Text** and **Table S1**). This periodic refresh of the host strain prevented undesired genomic adaptations from accumulating, forcing library genes to be the sole source of adaptation. On average, evolution campaigns were continued for ∼12 passages under selective conditions. The regularity of this workflow enabled us to complete dozens of evolution campaigns in parallel in under one month, spanning multiple selection pressures (see next section), error-prone DNAPs, starting ORF libraries, and one or two replicates per **EVO** experiment.

At the end of each **EVO** (∼100 generations), the population of adapted genes was isolated by **PCR** and integrated into a fluorescently marked selection strain for expression from a non-hypermutating genomic locus (**INT**_genome_). The non-mutating population was then competed against a different fluorescently marked strain encoding a nonsense gene (COMPETITION_population/NS_ or **COMP**_pop/NS_) to confirm that the evolved gene population was responsible for adaptation. If so, competition of the evolved gene population, which we empirically observed was always descended from a single gene from the initial gene library by the endpoints of evolution experiments, was repeated against the unevolved WT parent gene (**COMP**_pop/WT_) to assess whether the population of evolved sequences had higher average fitness than their unmutated ancestor. Individual mutant clones were then sampled, sequenced, and subjected to competitive growth experiments to confirm their individual fitness values against control strains expressing nonsense (**COMP**_mut/NS_) or corresponding WT genes (**COMP**_mut/WT_). Finally, custom experiments on evolved genes were carried out to dissect mechanisms by which they achieved their new functions.

### Design and implementation of selection pressures

To observe the emergence and evolution of diverse new gene functions, we chose multiple selection pressures for ORACLE experiments. The design of selection pressures employed in this study is as follows (**Fig. 1A**).

***Ev-Tol*** (**Tol**erance): The first set of experiments we carried out relied on non-optimal growth conditions imposed by the composition of the growth media. In these evolution campaigns, growth was limited either by deprivation of essential vitamins or the toxicity caused by elevated concentrations of certain ions in the media. This included growth media lacking vitamin B5 (pantothenate) (*44*) and high concentrations of manganese, lithium, and copper. The aim of these experiments was to observe the emergence and evolution of diverse genes conferring tolerance to toxic or suboptimal growth environments like what might be commonly encountered in nature.

***Ev-TF*** (**T**ranscription **F**actor): In this set of ORACLE experiments, we asked whether we could discover and evolve genes to obtain novel transcription factor activity, as changes in gene transcription are a common evolutionary challenge in nature. To link growth to transcription factor function, we placed *ADE2*, which encodes an enzyme responsible for a key step in adenine biosynthesis (*45*) and is essential for growth in media lacking adenine, under the control of the phosphate responsive *S. cerevisiae PHO5* promoter (pPHO5). In normal growth conditions, pPHO5 is covered by nucleosomes positioned to block access to the TATA box and other cis-regulatory elements, resulting in low transcription (*46*, *47*). Upon phosphate starvation, the Pho4 transcription factor can enter the nucleus, evict the nucleosomes and recruit transcriptional machinery (*47*). Our evolution experiments were carried out in high phosphate adenine-deficient media conditions, forcing ORACLE to discover and evolve genes that mimic the Pho4 transcription factor or otherwise activate its program.

***Ev-Deg*** (**Deg**ron): A major mechanism by which evolution changes the phenotypes of cells is by modifying the steady state level of proteins in the proteome. In this category of ORACLE experiments, we wanted to test whether we could evolve new genes that inhibited the ubiquitin-mediated degradation of a conditionally essential enzyme. We again used *ADE2* as the selectable marker to create a titratable growth-based selection. However, rather than controlling *ADE2*’s transcription, we destabilized the Ade2 protein using the ubiquitin–proteasome degradation pathway in yeast by appending N-terminal ubiquitin-based degradation tags to Ade2. In such systems, an N-terminal ubiquitin fused to Ade2 is first cleaved by deubiquitinases (DUBs), exposing a defined N-terminal residue on Ade2 (*48*). Then, the N-terminal amino acid dictates the half-life of Ade2 by the so-called N-end rule, which acts through the recruitment of ubiquitin ligases to Ade2 to mark the protein for degradation (*48–50*).

We implemented two selection configurations in this category. For the first, *ADE2* was placed under control of a weak promoter (pPOP6) and fused to a medium-strength N-terminal ubiquitin–tyrosine degradation tag (UbiY), yielding the pPOP6-UbiY-*ADE2* selection (*48*, *51*). In the second configuration, *ADE2* was driven by the strong pTDH3 promoter and fused to a more strongly destabilizing ubiquitin–arginine tag (UbiR), forming the pTDH3-UbiR-*ADE2* selection. For simplicity, these two configurations are referred to as ***Ev-Deg-Low*** (pPOP6-UbiY-*ADE2*) and ***Ev-Deg-High*** (pTDH3-UbiR-*ADE2*). In both setups, the translationally fused N-terminal ubiquitin is hydrolyzed by DUBs to generate an Ade2 with a defined N-terminal amino acid (tyrosine or arginine). The half-life of the resulting Ade2 is then controlled by the N-terminal amino acid’s propensity to be a client of the ubiquitin ligase, Ubr1, which recognizes hydrophilic N-terminal residues in its type I site or hydrophobic residues in its type II site and has additional activity for misfolded substrates (*52*). In **EVO** experiments, we selected for high Ade2 abundance through continuous passaging in media lacking adenine. Adaptation could occur through the inhibition of Ubr1 recognition of the N-terminal residue, by perturbing any number of associated processes that mediate degradation through N-terminal residues, by inhibiting the initial hydrolysis of the translationally fused ubiquitin that generates the defined N-terminus, by clogging protein degradation in general, etc.

### Overview of evolution results

ORACLE experiments successfully demonstrated clear examples of adaptation across all categories of selections, providing a spectrum of biomolecular innovations that revealed progressively more profound traversals of sequence-function space. At the simplest level, we observed adaptation through the rediscovery and evolution of known gene functions, representing evolutionary optimization along already established functional trajectories. Four examples of successes in this category are described. Moving beyond optimization of the primary functions of genes, we also observed adaptation through the discovery and evolution of unexpected gene functions. In these instances, evolution successfully co-opted and amplified promiscuous or latent weak gene activities, essentially repurposing an existing sequence for a novel primary role similar to the IAD model of neofunctionalization (*5*, *15*). We present and characterize three such examples. Ultimately, the most profound demonstration of gene evolution was adaptation through the *de novo* emergence of a function. We describe seven such examples, including the generation of a functional peptide sequence through a frameshift mutation, the emergence of specific transcription factor activity from a cryptic prophage gene, and the origination of several modulators of ubiquitin-mediated protein degradation pathways from uncharacterized proteins. **Table 1** provides a summary of these key findings, which are described in detail as a series of accounts, while **Table S1** summarizes all experiments done in this study and their outcomes. **Fig. 2-5** summarize key evolutionary outcomes and their characterization from ***Ev-Tol***, ***Ev-TF***, ***Ev-Deg-*Low**, and **Ev*-Deg-High*** evolution experiments, respectively.

**Fig. 2.**
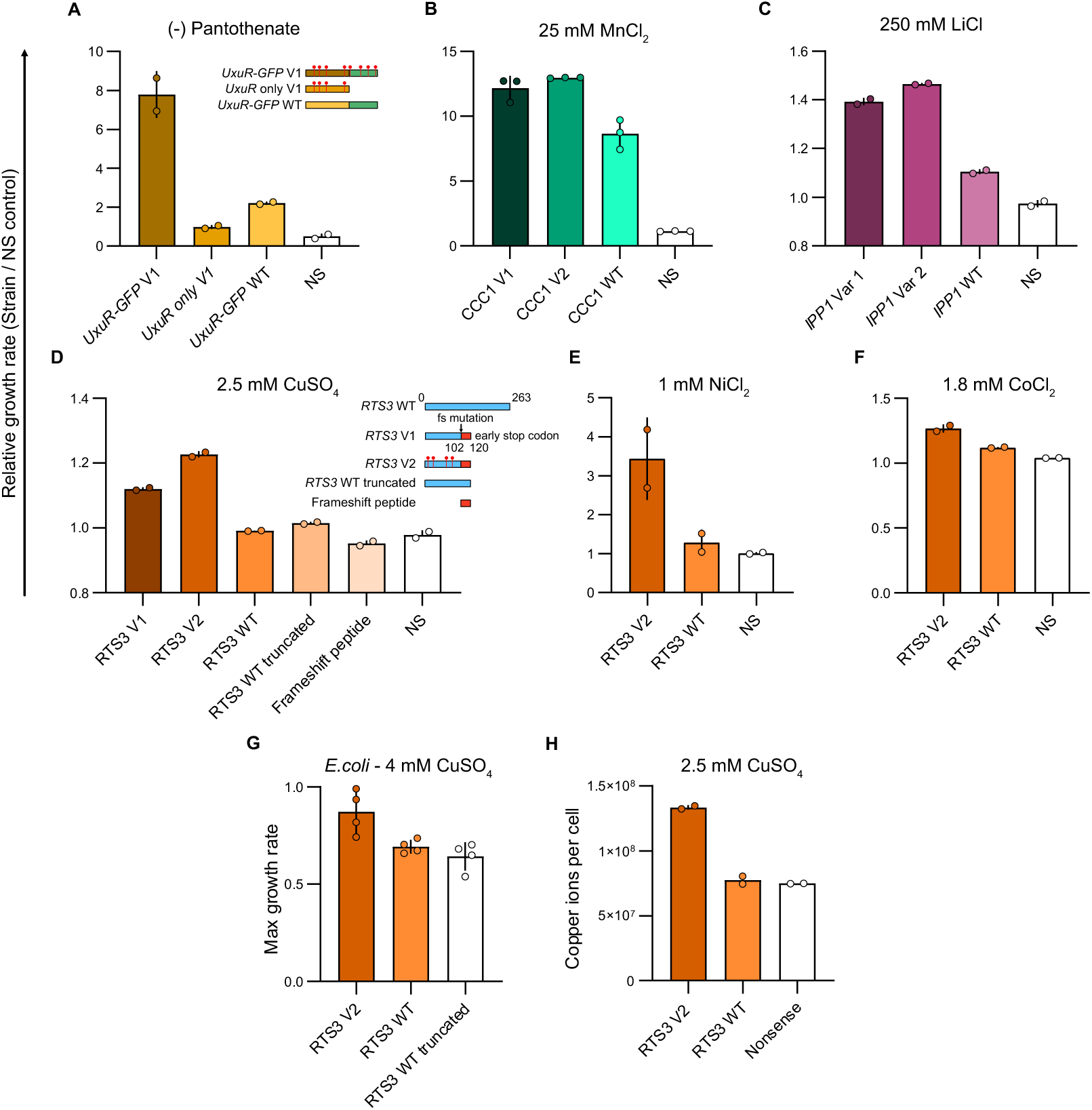
*Ev-Tol* evolution outcomes. **(A)** Relative growth rates against NS control for evolved *UxuR-GFP Variant 1*, WT *UxuR-GFP*, and *UxuR Variant 1* without GFP in media lacking pantothenate. **(B)** Relative growth rates against NS control of two *CCC1* variants and WT *CCC1* demonstrating fitness improvement conferred by evolved variants in media containing toxic manganese concentrations (25 mM, Manganese II chloride). **(C)** Relative growth rates against NS control of two *IPP1* variants and WT *IPP1* demonstrating fitness improvement conferred by evolved variants in media containing a toxic concentration of LiCl (250 mM). **(D-H)** A frameshift mutation in *RTS3* increased metal tolerance in yeast with *de novo* creation of a metal binding sequence. **(D)** Relative growth rates against NS control for evolved *RTS3* variants, WT *RTS3*, a WT *RTS3* artificially truncated to the same length as evolved *RTS3* variants, and the isolated 18 AA peptide resulting from the frameshift in *RTS3 Variant 2*, all in 2.5 mM CuSO_4_. The evolved *RTS3 Variant 2* also conferred resistance to nickel **(E)** and cobalt **(F)**, whereas WT *RTS3* did not. **(G)** The evolved *RTS3 Variant 2* increased copper tolerance in *E. coli*, while the WT *RTS3* did not. Max growth rate was extracted from a 24-hour growth curve at 37 °C, and the doubling time was calculated as described in **Materials and Methods** for two bioreplicates and two technical replicates. Error bars indicate standard deviation. Growth rates here are in units per hr. **(H)** *RTS3 Variant 2* overexpression increased the amount of intracellular copper in high copper conditions, as measured by ICP-MS, indicating that the evolved variant likely causes sequestration of copper. For (A-F), relative growth rates of NS against NS were determined as controls to ensure no difference (*i.e.*, relative growth rates of 1.0). For (A-F), competition experiments were done in 2-4 replicates with individual datapoints plotted around each bar depicting the mean. Error bars indicate standard deviation.

**Table 1.** Summary of 10 key ORACLE experiments carried out along with 19 evolved gene variants representing evolutionary outcomes demonstrating clear enhanced or new gene functions (see **Table S2** for sequences).

| Experiment # | Gene library | Selection | Gene identified | Nonsynonymous mutations | Mechanistic hypothesis |
| --- | --- | --- | --- | --- | --- |
| 1 | Yeast | *Ev-Tol-Pan* | *FMS1 Variant1* | V10A, K428R, I481V, A488V | Enhanced pantothenate synthesis |
| 2 | *E. coli*-GFP | *Ev-Tol-Pan* | *UxuR-GFP Variant1* | A32T, R67H, R150C, G215S, *GFP* Y343H, H358Y, K415E, I429T | Induction of low expression yeast pantothenate synthesis genes |
| 3 | Yeast | *Ev-Tol-Mn* | *CCC1 Variant1* | S68P, F92L, T274A, A313T | Improved activity for Mn import into vacuoles, overexpression better tolerated |
| 3 | Yeast | *Ev-Tol-Mn* | *CCC1 Variant2* | S68P, F256S, T274A, A313T | Improved activity for Mn import into vacuoles, overexpression better tolerated |
| 4 | Yeast | *Ev-Tol-Li* | *IPP1 Variant1* | K11D, I69V, F190L, K199R, K228E, T251A | Lithium binding and sequestration |
| 4 | Yeast | *Ev-Tol-Li* | *IPP1 Variant2* | K11D, I69V, P167T, A232V, T251A | Lithium binding and sequestration |
| 5 | Yeast | *Ev-Tol-Cu* | *RTS3 Variant1* | c.307delC (frameshift) | Copper binding and sequestration in nontoxic form |
| 5 | Yeast | *Ev-Tol-Cu* | *RTS3 Variant2* | V8A, A19T, V39I, K74R, c.307delC (frameshift) | Copper binding and sequestration in nontoxic form |
| 6 | *E. coli*-GFP | *Ev-TF* | *YdfC-GFP Variant1* | D12V, A54V, *GFP* R91H, R165H, H173Y, V204I, R260H, I263T, A298V, V311I, L313H | Transcription factor / chromatin regulating activity |
| 7 | *E. coli* | *Ev-TF* | *YeiG Variant1* | E27G, G89R, A189V, P268S | Transcription factor activity |
| 8 | Yeast | *Ev-Deg-Low* | *PET18 Variant1* | Y41H, F49S, S196N | Stabilization of N' terminal Ubi-X by direct binding to Ubi-X |
| 8 | Yeast | *Ev-Deg-Low* | *PET18 Variant2* | K74E, A142T, E143D, R148K | Stabilization of N' terminal Ubi-X by direct binding to Ubi-X |
| 9 | Yeast | *Ev-Deg-High* | *RPS31 Variant1* | R74G | Delay cleavage and processing of Ubiquitin fusion proteins by DUBs |
| 9 | Yeast | *Ev-Deg-High* | *ROQ1 Variant1* | L2S, Y88H | Inhibit Ubr1 activity toward N-degron substrates |
| 9 | Yeast | *Ev-Deg-High* | *ROQ1 Variant2* | I65T | Inhibit Ubr1 activity toward N-degron substrates |
| 10 | *E. coli* | *Ev-Deg-High* | *YjiE Variant1* | F143S | Global degradation inhibition |
| 10 | *E. coli* | *Ev-Deg-High* | *Aat Variant1* | L19P, I62V | Transfer of longer-lived amino acids to N' terminal arginine and lysine |
| 10 | *E. coli* | *Ev-Deg-High* | *YdaG Variant1* | A17T, V18M, C30Y, V38M | Specific inhibition of Ubr1 |
| 10 | *E. coli* | *Ev-Deg-High* | *SecB Variant1* | A14T, A41T, Q49L, D59N, V80I, A145T | Chaperone activity for *UbiR-Ade2* |

### Adaptation through rediscovery and evolution of known gene function

***Ev-Tol-Pan:* EVO** experiments from the yeast ORF library exposed to evolution in pantothenate dropout yielded the fixation of lineages descended from two genes, *FMS1* and *SPE4*, which is consistent with their prior implication in an alternative pantothenate synthesis pathway (*53*). Therefore, the outcome of this experiment is the rediscovery of a known pathway that alleviates pantothenate deprivation. Nevertheless, **EVO** was able to improve the enriched genes. For example, we enriched a superior *FMS1* clone containing four mutations (V10A, K428R, I481V, A488V) that rendered it superior to WT *FMS1* in conferring tolerance to pantothenate dropout (**Fig. S7**), as confirmed by **COMP**_mut/WT_. In fact, the *FMS1* V10A, K428R, I481V, A488V (*FMS1 Variant 1*) mutant was able to support growth in pantothenate dropout media at a rate comparable to that in complete media. Notably, there was no fitness cost for this mutant in pantothenate-rich media, indicating minimal metabolic burden (**Fig. S7F**).

***Ev-Tol-Mn:*** For manganese tolerance **EVO** experiments, sequencing revealed the fixation of lineages descended from a yeast gene called *CCC1*. *CCC1* encodes a vacuolar transporter whose overexpression is known to mediate tolerance to toxic Mn^2+^ concentrations by import into the vacuole (*54*). Therefore, it was not surprising that our **EVO** experiments enriched *CCC1* lineages. Nevertheless, the actual evolved *CCC1*’s mutants, specifically variants carrying mutations S68P, T274A, A313T, and an additional F92L (*Variant 1)* or F256S (*Variant 2*) mutation, were superior to WT *CCC1* in two ways. First, these variants enhanced manganese tolerance (**Fig. 2B** and **Fig. S8**) more than WT *CCC1*, as determined by **COMP**_mut/WT_. Second, in normal conditions lacking manganese, overexpression of WT *CCC1* conferred a substantial growth defect whereas overexpressing the evolved *CCC1* variants carried less fitness defect (**Fig. S8C**). Therefore, *CCC1* evolved an important new overall function: greater activity and specificity in transporting Mn^2+^.

***Ev-Deg-High***: **EVO** from yeast ORF libraries under ***Ev-Deg-High*** selections yielded successful lineages descended from a recently discovered Ubr1 modulator, Roq1 (*52*, *55*). Roq1 is known to alter the substrate specificity of Ubr1 away from hydrophilic N-terminal amino acids towards misfolded clients by acting as a pseudosubstrate that allosterically regulates Ubr1 with multiple cooperating motifs (*52*, *55*). Given this known function, it is unsurprising that *ROQ1* emerged from evolution. However, it is notable that the two evolved *ROQ1* gene variants we isolated, containing nonsynonymous mutations L2S, Y88H (*Variant 1*) and I65T (*Variant 2*), showed increased fitness over WT *ROQ1* in **COMP**_mut/WT_ assays (**Fig. 5A** and **Fig. S9**). This suggests that the mutations increase Roq1 activity. Interestingly, I65K and I65A mutations were previously identified to make *ROQ1* less active (*55*), while I65T was an activity-increasing mutation our evolution experiment found.

To further characterize our *ROQ1* variants on Ubr1-driven degradation, we established a set of luciferase-based reporter assays in which a luciferase was fused downstream of various UbiX tags to generate luciferases containing X as the N-terminal residue after ubiquitin is removed by DUBs. Ubr1 recognizes these N-termini using its type I site for positively charged N-terminal residues (*e.g.*, R and K) or type II site for bulky hydrophobic residues (*e.g.*, Y). Hence, a general Ubr1 or degradation inhibitor should stabilize UbiR-, UbiK- and UbiY-Luciferases, while a specific type I site modulator should only stabilize UbiR- and UbiK-Luciferases, having no effect on UbiY-Luciferase. UbiM-Luciferase was included as a control since M is a stabilizing N-terminal residue that is not a client for Ubr1. Additionally, PEST-mediated degradation of GFP was also measured, since PEST-tagged proteins are recognized and ubiquitinated by a different machinery (SCF complex) and are therefore Ubr1 independent (*56*). We reasoned that inhibitors of global protein degradation (*e.g.*, proteasome inhibitors) would stabilize both UbiX-Luciferase and the GFP-PEST reporter while any Ubr1-specific modulators should have no effect on the GFP-PEST reporter. Lastly, Ubr1 is known to enhance degradation of certain misfolded proteins through a putative separate substrate-binding site distinct from its N-degron recognition domains. We therefore additionally measured the effect of each evolved hit on degradation of a known misfolded substrate of Ubr1, Rtn1Pho8* (*52*), using a Rtn1Pho8*-GFP reporter.

In luciferase reporter assays, both the evolved *ROQ1* variants were found to more strongly prevent the degradation of UbiR-tagged and UbiK-tagged luciferase compared to WT *ROQ1* (**Fig. 5B**). This confirms the evolution of a more potent Roq1. In the assays for PEST-mediated degradation and misfolded protein degradation, the evolved Roq1s and the WT Roq1 redirected Ubr1 comparably (**Fig. 5C**). These data demonstrate that the evolved variants selectively enhance Roq1’s capacity to block Ubr1’s canonical N-degron pathway without modulating its allosteric promotion of misfolded protein degradation.

**EVO** from the *E. coli* ORF library under ***Ev-Deg-High*** selections also yielded an example where a known primary gene function was the source of adaptation. Specifically, we observed the fixation of lineages derived from the *E. coli aat* gene. This gene encodes a leucyl/phenylalanyl-tRNA-protein transferase, which attaches leucine and phenylalanine to N-terminal arginines and lysines (*49*, *57*). The likely mechanism through which Aat likely confers fitness is obvious: leucine is made to be the new N-terminal amino acid of Ade2, imbuing Ade2 with a longer half-life given the preferences of Ubr1 (*49*, *52*). However, **EVO** did not simply enrich WT *aat*, but rather generated an improved variant, with mutations L19P and I62V. The activity improvement over WT *aat*, as determined through fitness by **COMP**_mut/WT_, was recapitulated in the luciferase reporter assays as well (**Fig. 5A-B** and **Fig. S10**).

### Adaptation through discovery and evolution of unexpected gene function

***Ev-Tol-Pan:*** Evolution in pantothenate dropout from an *E. coli* ORF library where each ORF was fused to GFP (*42*, *43*) yielded a mutant transcriptional repressor-GFP fusion, UxuR-GFP, that dramatically increased fitness compared to WT UxuR-GFP, which only modestly increased fitness as determined by **COMP**_mut/NS_ and **COMP**_WT/NS_ (**Fig. 2A** and **Fig. S7G**). The evolved UxuR-GFP *Variant 1*, containing A32T, R67H, R150C, G215S, Y343H, H358Y, K415E, I429T mutations, required the GFP portion of the fusion for its functionality, possibly due to the stability that GFP may confer (**Fig. 2A** and **Fig. S7H**). Although it is not obvious how the evolved UxuR-GFP might mechanistically achieve tolerance to pantothenate dropout, this experiment demonstrates that a non-native gene can have unexpected activity, allowing it to enter the view of selection for improvement in a new organism. This highlights the potential role of horizontal gene transfer in the evolutionary origins of unexpected traits.

***Ev-Tol-Li:*** The yeast ORF library and the *E. coli* ORF library provided successful adaptation under inhibitory lithium chloride conditions, as assayed by **COMP**_pop/NS_. In these cases, lineages derived from two inorganic pyrophosphatase genes, yeast *IPP1* and *E. coli ppa* (**Fig. 2C** and **Fig. S11**), fixed. Despite their low overall sequence identity (22–27%), the active sites of these two genes are highly conserved (82–94% identity) (*58*) and contain multiple Mg^2+^ binding sites (*59*). It is plausible that these sites also confer lithium binding activity, explaining their fitness benefit in ***Ev-Tol-Li*** in retrospect. **COMP**_pop/WT_ assays showed substantial adaptation beyond the ancestral WT *IPP1* for the yeast ORF library **EVO** outcomes. We therefore sampled two variants of *IPP1*s at the end of **EVO** for further testing. These two variants shared mutations K11D, I69V, T251A, but had additional unshared mutations (*Variant 1*: plus F190L, K199R, K228E; *Variant 2*: plus P167T, A232V). Both variants indeed conferred high lithium tolerance exceeding the activity of the WT *IPP1* sequence (**Fig. 2C** and **Fig. S11C**). Since the WT *IPP1* sequence still conferred some tolerance (**Fig. 2C** and **Fig. S11**), this **EVO** experiment can be described as a case where the WT *IPP1* gene first enriched from the library through its latent weak lithium tolerance activity and then evolved through a multi-mutational pathway towards high lithium tolerance, further highlighting how gene functions can arise from unexpected sources.

***Ev-TF:*** In **EVO** experiments for driving transcription from the pPHO5 promoter, we discovered a surprising result wherein the *YeiG* gene from the *E. coli* ORF library evolved successfully. WT *YeiG* showed modest activity in inducing expression from pPHO5, while evolved mutants collected at the end of **EVO** showed high activity (**Fig. 3A** and **Fig. S12**). In *E. coli*, *YeiG* encodes a serine hydrolase with primary activity on the substrate, S-formylglutathione (*60*). To the best of our knowledge, no transcription factor activity for *YeiG* has been described. Therefore, this is another case of unexpected gene activity becoming the start of an adaptive walk.

**Fig. 3.**
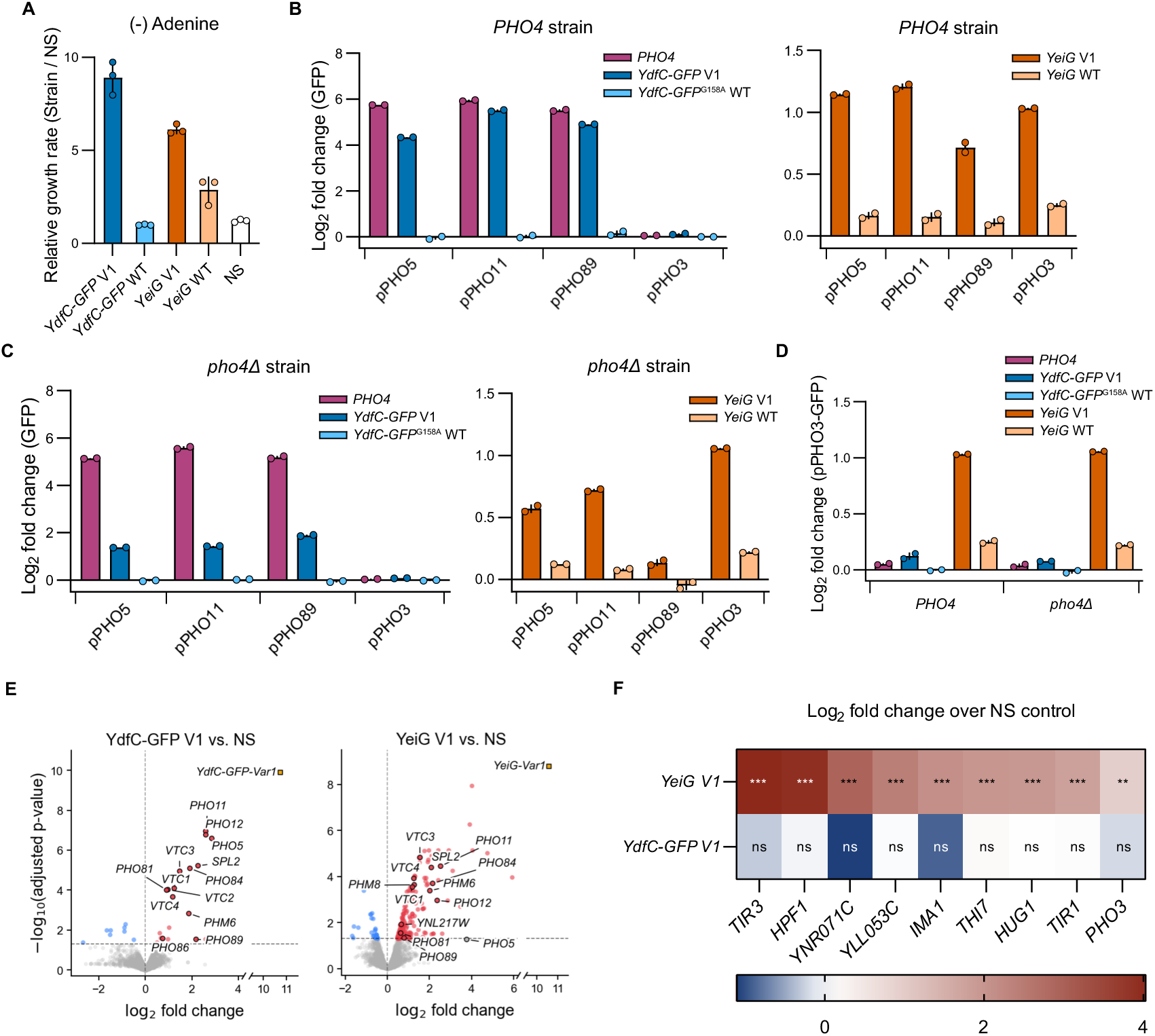
*Ev-TF* evolution outcomes. **(A)** Relative growth rates against NS control for *YdfC-GFP Variant 1*, WT *YdfC-GFP*, *YeiG Variant 1*, and WT *YeiG* in the pPHO5-*ADE2* selection strains in the absence of adenine. Relative growth rate of NS against NS was determined as a control to ensure no difference (*i.e.*, relative growth rates of 1.0). Competition experiments were done in triplicate with individual datapoints plotted around each bar depicting the mean. Error bars indicate standard deviation. **(B-C)** Induction of phosphate responsive promoters by various genes in *PHO4* and *pho4*Δ strains. GFP was placed under the control of each promoter, and its fluorescence was measured in the presence of each gene alongside fluorescence in the presence of a nonsense control gene’s expression. Data plotted is the log_2_ of the fold change in GFP geomean fluorescence induced by expression of each gene compared to that induced by expression of a nonsense control gene. Experiments were done in duplicate with individual datapoints plotted around each bar depicting the mean log_2_ of fold change. Error bars indicate standard deviation. Note that we introduced a G158A mutation into WT *YdfC-GFP* for these assays to negate any confounding fluorescence from the gene, which was not necessary for the evolved *YdfC-GFP* variants, since they already contain mutations in the GFP domain that rendered it nonfluorescent. **(D)** Comparison of induction for natively downregulated *PHO3* promoter with or without the genomic *PHO4*, demonstrating distinct induction profile caused by overexpression of *YeiG Variant 1*. **(E)** The evolved *YeiG Variant 1* and *YdfC-GFP Variant 1* upregulated the expression of many canonical Pho4-responsive genes, with *YdfC-GFP Variant 1* conferring a much more specific upregulation of the canonical Pho4-responsive genes. Volcano plots of RNA-seq data show the distribution of fold-change and statistical significance for upregulated and downregulated genes. Gene expression changes were calculated by comparing a strain overexpressing the evolved gene (gold square) to a nonsense control strain. Pho4-responsive genes are represented as dots with dark borders. RNA-seq for *YeiG Variant 1* and *YdfC-GFP Variant 1* was performed with 2 and 3 biological replicates, respectively. **(F)** Heat map showing the gene expression fold change for a set of Pho4*-*independent genes, including *PHO3*, demonstrating that *YeiG Variant 1* activated transcription of an array of genes not associated with the Pho4 regulon, while *YdfC Variant 1* did not. **p < 0.01, ***p < 0.001; ns, not significant (adjusted p-values). RNA-seq for *YeiG Variant 1* and *YdfC-GFP Variant 1* was performed with 2 and 3 biological replicates, respectively.

To dissect mechanism, we carried out a set of experiments aimed at characterizing the evolved *YeiG*s’ transcription factor activities. We built a series of yeast reporter strains where various phosphate response promoters, pPHO5, pPHO11, and pPHO89, as well as the constitutively active pPHO3 promoter (*61*), were cloned upstream of a GFP reporter gene and integrated into the genome. pPHO5, pPHO11, and pPHO89 are three canonical promoter targets of the Pho4 transcription factor, whereas the pPHO3 promoter drives constitutive expression that serves as a non-responsive control here. We verified that GFP expression reflected the expected profile in phosphate starvation conditions – induction at the pPHO5, pPHO11, and pPHO89 promoters, while expression from the pPHO3 promoter remained unchanged (**Fig. 3B** and **Fig. S12C**) (*61*). We then tested an evolved *YeiG* variant (*Variant 1*, containing mutations E27G, G89R, A189V, P268S) for activity in these reporter strains in normal conditions without phosphate starvation. We found that while WT *YeiG* effected a slight increase in expression from the inducible phosphate starvation response promoters, *YeiG Variant 1* drove much higher expression (**Fig. 3B** and **Fig. S12E**). To determine whether the evolved *YeiG* was inducing a response independently of the Pho4 transcription factor that normally occupies and activates pPHO5 under phosphate starvation, we repeated our reporter assay with yeast’s endogenous copy of *PHO4* knocked out. We found that the transcriptional activation strength of the evolved *YeiG Variant 1* was partially reduced in some cases, but not eliminated, in the *PHO4* deletion strain (**Fig. 3C** and **Fig. S12E**). Interestingly, expression from the pPHO3 promoter, which is unaltered by phosphate starvation or *PHO4* overexpression, was upregulated by the evolved *YeiG Variant 1* (**Fig. 3D** and **Fig. S12E**).

To further profile *YeiG Variant 1*’s function, we carried out RNA-seq experiments to compare the transcriptome of a strain expressing *YeiG Variant 1* with that of a control strain encoding a nonsense gene, all under non-selective (adenine-rich) conditions. This experiment revealed that overexpression of the evolved *YeiG* upregulates transcription of 13 classic Pho4 target genes, including *PHO5*, *PHO11*, and *PHO81* (**Fig. 3E** and **Fig. S13**) (*62*). However, *YeiG* also displayed a broader transcriptional activity beyond these canonical targets, upregulating several genes not typically associated with the Pho4-mediated response – such as *PHO3* (**Fig. 3F**). Taken together, our experiments are consistent with the hypothesis that *YeiG Variant 1* functions as a transcription factor similar to but more promiscuous than Pho4, driving expression of both the canonical phosphate regulon and a diverse set of non-canonical loci. Other mechanisms of action are also plausible given the broad expression profile changes induced. From a gene evolution perspective, this example of *YeiG* evolution may be a case where an unrelated moonlighting function of an enzyme begins its evolution of regulatory protein functionality.

### Adaptation through de novo emergence and evolution of gene function

***Ev-Tol-Cu:* EVO** experiments for copper tolerance from the yeast ORF library yielded the fixation of lineages descended from the *S. cerevisiae RTS3* gene. *RTS3* encodes a largely unstructured protein previously linked to caffeine tolerance (*63*). Strikingly, the evolved variants of *RTS3* that enriched after **EVO** contained a single nucleotide deletion, producing a frameshifted, truncated protein with 120 amino acids (instead of *RTS3*’s original 263 amino acid size) and 18 new amino acids prior to the stop codon. The last 18 amino acids of this evolutionary outcome can therefore be considered a *de novo* peptide. Furthermore, *RTS3* with the frameshift alone (*Variant 1*) or with additional mutations discovered by the end of **EVO** (*Variant 2*, including the frameshift plus V8A, A19T, V39I, and K74R) conferred copper tolerance, while the WT *RTS3* conferred none (**Fig. 2D** and **Fig. S14**). Thus, we can describe this **EVO** example as one in which a new gene function originated *de novo*.

We note several interesting observations about the frameshifted *RTS3*. First, simply truncating WT *RTS3* to create a 120 amino acid fragment was not sufficient to observe any copper tolerance, demonstrating that the 18 amino acid *de novo* peptide in the frameshifted *RTS3* was critical for function (**Fig. 2D**). Second, expression of only the 18 amino acid *de novo* peptide portion of the frameshifted *RTS3* showed no effect on copper tolerance (**Fig. 2D** and **Fig. S14B**), suggesting that the peptide appendage needs the remainder of *RTS3* for its discovered function, if only for stabilization, since short peptides are typically degraded. Third, the most active mutant, *Variant 2* with four additional substitution mutations, conferred higher copper tolerance than *Variant 1* (**Fig. 2D** and **Fig. S14C**). This is consistent with an evolutionary pathway where WT *RTS3*, with no detectable initial activity for copper resistance, randomly sampled a single nucleotide frameshift mutation to introduce an 18 amino acid *de novo* peptide appendage that originated weak copper tolerance, which was then improved by further evolution.

To investigate mechanism, *RTS3 Variant 2* was tested for its ability to confer tolerance to other divalent ions, such as nickel (**Fig. 2E** and **Fig. S14D**) and cobalt (**Fig. 2F** and **Fig. S14E**), using **COMP**_mut/NS_ assays. These assays revealed that *RTS3 Variant* 2’s function extended to the additional metals. Furthermore, we expressed the frameshifted *RTS3*s in *E. coli*, where they were also found to increase tolerance to copper (**Fig. 2G**).

The generality of contexts in which the evolved frameshifted *RTS3s* confer divalent metal ion tolerance suggests they may be acting to sequester ions directly. To explore this possibility, we measured intracellular copper concentrations using ICP-MS under high copper stress in yeast. Yeast strains expressing either WT *RTS3*, *RTS3 Variant 2*, or a nonsense gene control were passaged twice in media containing 2.5 mM CuSO_4_. We found that the intracellular copper content in cells was substantially elevated in the samples expressing *RTS3 Variant 2* compared to both WT *RTS3* and the nonsense control (**Fig. 2H**). We therefore presume that the copper tolerance is mediated by copper sequestration inside the cells, which may also include the nucleation of biomineralization or phase separation, limiting the potential of copper to bind sensitive targets. That *RTS3 Variant 2*’s copper tolerance function owes to a *de novo* peptide appendage supports previous literature suggesting the relative accessibility of metal-binding activity in *de novo* proteins (*64*).

***Ev-TF:*** In another example where a gene function both originated *de novo* and subsequently evolved, ORACLE yielded presumptive new transcription factors from a cryptic prophage gene. From the *E. coli* ORF library fused to GFP, we found that an ***Ev-TF*** campaign resulted in the fixation of lineages derived from a cryptic prophage peptide, YdfC from the Qin prophage (*65*). Specifically, we isolated a *YdfC-GFP* variant, *Variant 1*, containing mutations (D12V, A54V)-GFP(R91H, R165H, H173Y, V204I, R260H, I263T, A298V, V311I, L313H), which confers high fitness compared to the WT *YdfC-GFP* fusion (**Fig. 3A** and **Fig. S12A-B**). Notably, WT *YdfC-GFP* had no activity as determined by **COMP**_WT/NS_, making this an example where a gene with no activity discovered an innovation through new mutations and further evolution.

In reporter assays characterizing the ability of *YdfC-GFP Variant 1* to drive phosphate starvation response promoters, we observed that the evolved *YdfC-GFP* induced the same expression profile as phosphate starvation (**Fig. 3B** and **Fig. S12D)**. The resulting expression profile closely mirrored the effect of overexpressing the endogenous Pho4 transcription factor that mediates the phosphate starvation program. In contrast, WT *YdfC-GFP* (containing a synthetically installed G158A mutation in GFP that makes it nonfluorescent so as not to confound the fluorescent readout from the reporter assay) (*66*) did not induce any expression of the phosphate starvation program (**Fig. 3B** and **Fig. S12D**). Furthermore, in a *PHO4* deletion strain, the evolved *YdfC-GFP Variant 1* still drove expression of the phosphate starvation program, albeit at a reduced level (**Fig. 3C** and **Fig. S12D**). This is consistent with the *YdfC-GFP Variant 1* acting as a transcription factor with a similar promoter binding and nucleosome eviction profile as Pho4.

RNA-seq experiments were consistent with this hypothesis, showing highly specific upregulation of 13 canonical Pho4 target genes (**Fig. 3E** and **Fig. S13**) (*62*). Furthermore, the expression of *PHO3* and other Pho4-independent genes were unaffected by *YdfC-GFP Variant 1* (**Fig. 3F**). Therefore, unlike the evolved *YeiG* outcome (see above) that effected a broad transcriptional profile change, *YdfC-GFP Variant 1* faithfully recapitulated the specific transcriptional program of Pho4. In summary, ORACLE was able to originate and evolve a new gene function that replicates Pho4 transcription factor activity from an *E. coli* cryptic prophage gene-GFP fusion with no known function and no initial activity.

***Ev-Deg-Low***: In ORACLE experiments under ***Ev-Deg-Low*** conditions starting from the yeast ORF library, we observed clear adaptation from one gene, *PET18*. This gene has previously only been studied for potential activity in a thiamine salvage pathway due to its homology to *THI20*, but in that study, no activity could be identified (*67*). **EVO** in a selection strain that replicates p1 with Trixy and in a selection strain that replicates p1 with BadBoy3 independently resulted in fixation of *PET18*-derived mutants with high fitness – *Variant 1* containing Y41H, F49S, S196N and *Variant 2* containing K74E, A142T, E143D, R148K, respectively. Notably, no mutations were shared between the two variants. Since the WT *PET18* showed undetectable activity in the ***Ev-Deg-Low*** selection context in **COMP**_WT/NS_ assays (**Fig. 4A** and **Fig. S15**), this is another example where a gene function was both discovered *de novo* and subsequently evolved through ORACLE.

**Fig. 4.**
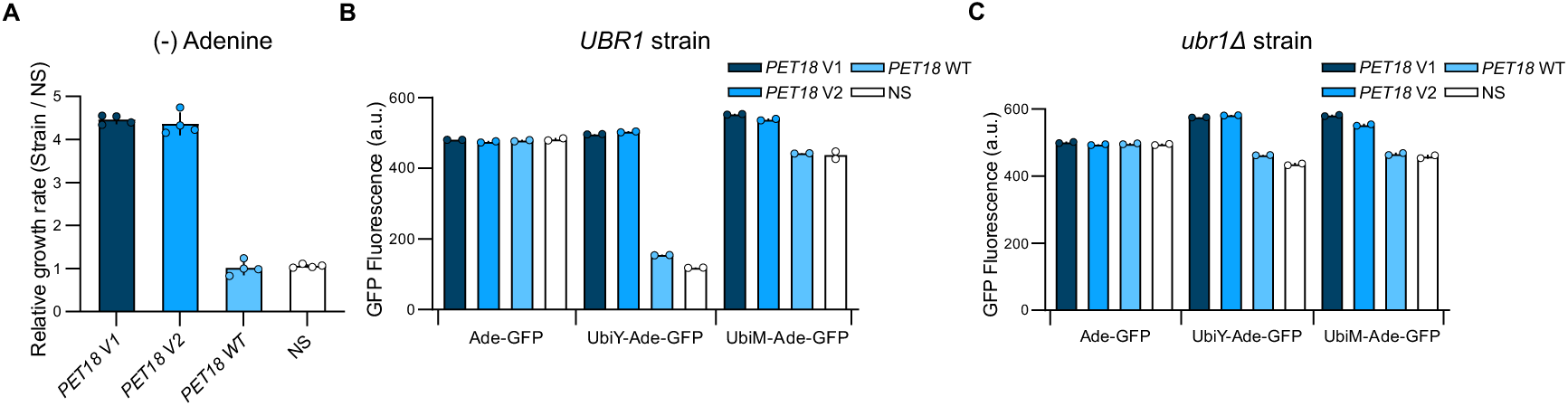
*Ev-Deg-Low* evolution outcomes. **(A)** Relative growth rates against NS control for *PET18 Variant 1*, *PET18 Variant 2*, and WT *PET18* in the pPOP6-UbiY-*ADE2* selection strains and media lacking adenine. Relative growth rate of NS against NS was determined as a control to ensure no difference (*i.e.*, relative growth rates of 1.0). Experiments were done in 4 replicates with individual datapoints plotted around each bar depicting the mean. Error bars indicate standard deviation. **(B-C)** Effect of overexpression of *PET18* variants and WT *PET18* on Ade2-GFP protein levels when expressed from pPOP6-*ADE2*-GFP, pPOP6-UbiY-*ADE2*-GFP, and pPOP6-UbiM-*ADE2*-GFP reporters in cells with and without *UBR1*. Data reported is average geomean GFP fluorescence measured in biological duplicate, with individual datapoints plotted around each bar. Error bars indicate standard deviation.

The two independently evolved *PET18* mutants, *Variant 1* and *Variant 2*, were subjected to further characterization. Recall that in the ***Ev-Deg*** experiments, Ade2 is expressed as a UbiX-Ade2 fusion, which is first processed by DUBs to hydrolyze Ubi and reveal X as the N-terminal amino acid of Ade2. Depending on the identity of X, the X-Ade2 protein is then a substrate for Ubr1, which ubiquitinylates Ade2 for degradation. Since we selected on the activity of Ade2 (using adenine dropout conditions), mechanisms by which evolved genes could achieve high fitness presumably included upregulation of UbiX-Ade2’s transcription, inhibition of the initial hydrolysis of Ubi by DUBs, inhibition of Ubr1’s engagement of X-Ade2 as a client, general inhibition of ubiquitin-mediated protein degradation, etc. We carried out a series of experiments to narrow down the mechanisms by which evolved *PET18* variants might be acting.

To measure the effects of overexpressing the evolved *PET18* variants, we developed a fluorescent reporter assay by fusing GFP to Ade2. As shown in **Fig. 4B**, the evolved variants protected Ade2-GFP from degradation when UbiY was attached (pPOP6-UbiY-*ADE2*-GFP) but had no effect on the pPOP6- *ADE2*-GFP reporter, suggesting that *PET18* variants do not function by simply increasing transcription from pPOP6. The effects were not specific for UbiY, as they reproduced with UbiF attached to Ade2-GFP (**Fig. S15B**), and slight increases in fluorescence could also be seen with UbiM N-terminally fused to Ade2-GFP (**Fig. 4B** and **Fig. S15A**). Consistent with this, the evolved *PET18*s also increased fitness in the context of other promoters driving the UbiY-*ADE2* construct (**Fig. S15B**) and increased fluorescence of a pPOP6-UbiX-GFP reporter lacking Ade2 in the fusion (**Fig. S15C**).

To determine whether *PET18* variants were inhibiting Ubr1 activity, we knocked out the *UBR1* gene and measured the effect of evolved *PET18* variants on various Ade2-GFP reporters. *UBR1*-deletion strains showed approximately the same level of fluorescence for the pPOP6-*ADE2*-GFP and pPOP6-UbiY-*ADE2*-GFP reporters in the absence of any *PET18*, confirming elimination of Ubr1 activity on the Y-Ade2-GFP fusion (**Fig. 4C**). We reasoned that if *PET18* variants were inhibiting Ubr1, their expression should have no effect on fluorescence from pPOP6-UbiY-*ADE2*-GFP in the *UBR1* deletion strain. Surprisingly, we observed that overexpression of the evolved *PET18* variants increased the fluorescence level of pPOP6-UbiY-*ADE2*-GFP relative to pPOP6-*ADE2*-GFP (**Fig. 4C**), suggesting a mechanism that involves an interaction with the Ubi domain of the UbiY-Ade2-GFP fusion. Consistent with this hypothesis, the growth rate of *UBR1* deletion strains containing the pPOP6-UbiX-*ADE2*-GFP cassette in the absence of adenine was also increased when evolved *PET18* variants were expressed (**Fig. S15D**).

Although it is possible that *PET18* variants also inhibit Ubr1 or general ubiquitin-mediated protein degradation, our most parsimonious hypothesis explaining these results is that the evolved *PET18*s act as chaperones that inhibit recognition of Ubi by DUBs, both stabilizing the UbiY-Ade2-GFP fusion and preventing it from hydrolysis into a Y-Ade2-GFP product that *Ubr1* recognizes. *PET18* shares structural homology to many aminopyrimidine aminohydrolases (*67*), and given its lack of known activity, may be a pseudogenized version of a thiaminase. The evolution of presumptive chaperone activity from a potential pseudogene with no known function is an example of how evolutionary innovation can spring from unexpected inert sources.

***Ev-Deg-High***: ORACLE also yielded several modulators of protein degradation in ***Ev-Deg-High* EVO** experiments, which involved the same UbiX-Ade2-based selection for Ade2 abundance as ***Ev-Deg-Low*** experiments, but where X is R (a stronger client for Ubr1 than Y) and where the expression of UbiR-Ade2 is driven by a strong promoter.

In addition to the *ROQ1* gene already described in a previous section, **EVO** from the yeast ORF library fixed a mutant *RPS31* gene containing only one mutation, R74G. Overexpression of the WT *RPS31* gene produced no detectable fitness benefit in **COMP**_WT/NS_ assays and no increase in luciferase levels in UbiX-Luciferase assays, while the evolved *RPS31* R74G mutant (*Variant 1*) showed substantial activity in both (**Fig. 5A-B** and **Fig. S9A-B**). Interestingly, when *RPS31 Variant 1* was tested for its effect on global protein degradation using the reporter assays where GFP was tagged with a PEST degron or a misfolded protein, no effect was observed (**Fig. 5C**), suggesting that *RPS31 Variant 1* does not broadly affect protein degradation. We note that *RPS31* encodes a ribosomal protein that is expressed with an N-terminal ubiquitin fusion. The first 76 residues represent ubiquitin, which is cleaved after Gly76 by DUBs to release the mature protein (*68*). Arg74 (R74), located in the flexible ubiquitin tail, is known to be crucial for this recognition and cleavage (*69*, *70*), and any substitution at Arg74 abolishes function (*71*). Therefore, the R74G mutation may be adaptive by turning *RPS31* into a DUB sponge that may partially prevent UbiX-protein fusions from being processed, allowing them to exist in their UbiX-protein fusion form without revealing the unstable N-terminus of their mature form. From a gene evolution perspective, this is an example where a single mutation originates a novel function.

**Fig. 5.**
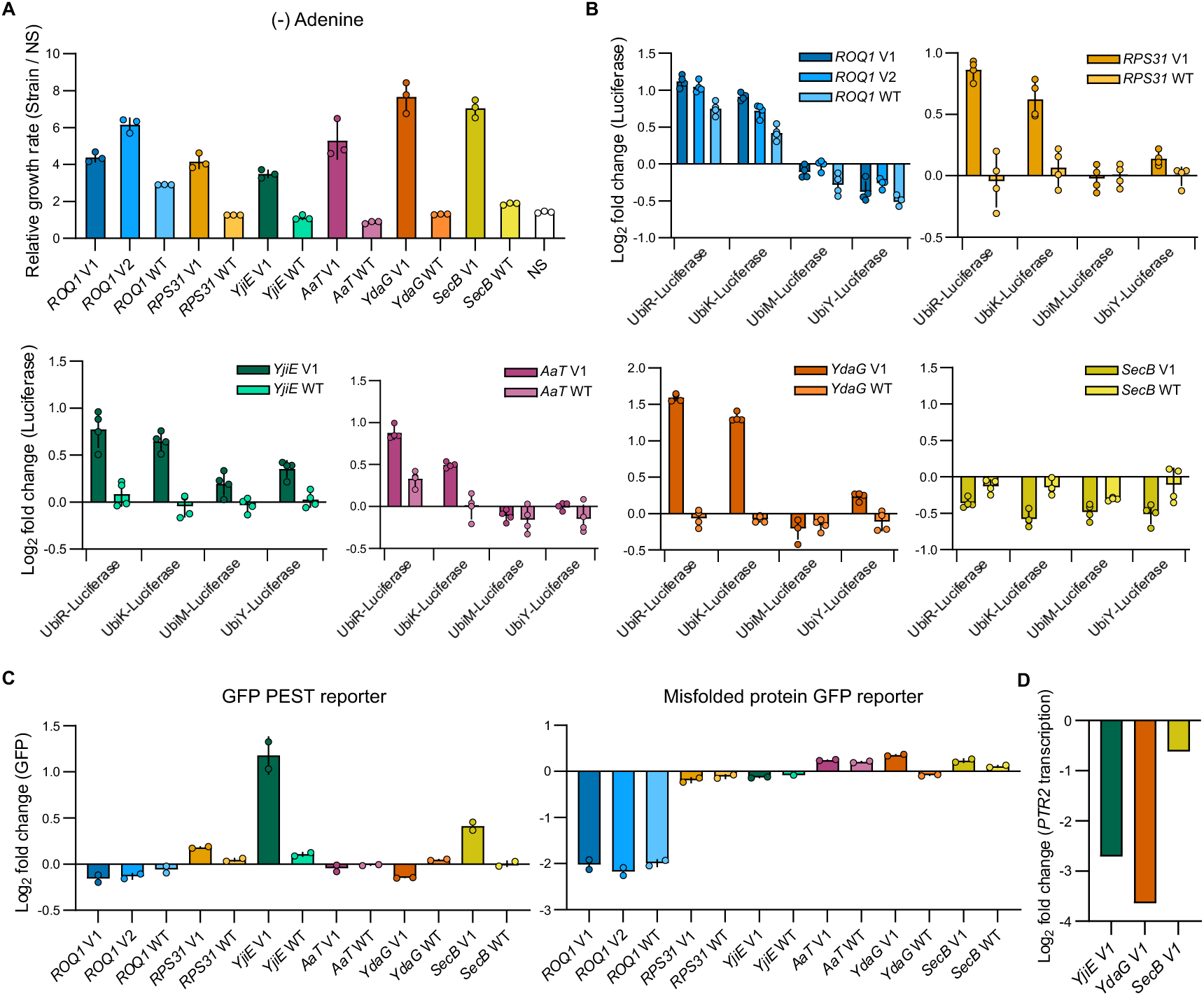
*Ev-Deg-High* evolution outcomes. **(A)** Relative growth rates against NS control for several evolved gene variants and their WT counterparts in the pTDH3-UbiR-*ADE2* selection strains and media lacking adenine. Relative growth rate of NS against NS was determined as a control to ensure no difference (*i.e.*, relative growth rates of 1.0). Experiments were done in 3 replicates with individual datapoints plotted around each bar depicting the mean. Error bars indicate standard deviation. **(B)** Luciferase reporter assay results measuring the stability of luciferase in UbiX-Luciferase fusions supports the stabilization activity of evolved variant genes and reveals more about their specific mechanisms. Activity of various UbiX-Luciferase fusions was assessed in strains overexpressing each of the evolved gene variants. Luciferase signal was measured in biological duplicate, and technical duplicate, and the log_2_ of fold change in signal over that of a nonsense control transformed alongside is reported. Individual datapoints are plotted around each bar depicting the mean. Error bars indicate standard deviation. Except for *SecB* genes, evolved variants of all tested genes increased levels of the Type I Ubr1 substrates (UbiR-and UbiK-Luciferase), with a subset also stabilizing the Type II substrate (UbiY-Luciferase). Notably, WT *ROQ1* and WT *aat* showed some activity in stabilizing the reporter luciferase, albeit at weaker levels than their evolved variants, whereas WT variants of other genes had undetectable activity. **(C)** Off-target reporter activity results evaluating the impact of evolved variants on degradation mediated by a PEST degradation tag (driven by SCF complex) or degradation of a misfolded reporter (Rtn1Pho8*-GFP) known to be substrate of Ubr1. Data reported is the log_2_ of fold change in geomean GFP fluorescence conferred by test genes over that conferred by expression of a nonsense control gene carried out alongside, measured in biological duplicate. Error bars indicate standard deviation. Evolved *YjiE Variant 1* decreased degradation of GFP-PEST indicating a decrease in global degradation. *ROQ1* variants were expected to redirect Ubr1 to increase degradation of the cytoplasmic misfolded reporter, as observed. *SecB Variant 1* slightly decreased the degradation of GFP-PEST. Other genes did not substantially impact degradation mediated by a PEST degradation tag or degradation of the misfolded reporter. **(D)** Log_2_ of *PTR2* transcription fold change for strains expressing the evolved variants of *YdaG*, *YjiE*, or *SecB* over a nonsense control strain, as obtained from RNA-seq data compiled from 3 replicates. Evolved *YdaG* and *YjiE* act as inhibitors of *Ubr1* and global degradation, respectively, which likely leads to *Cup9* accumulation and therefore strong *PTR2* downregulation. Meanwhile, *SecB Variant1* fails to show comparable *PTR2* downregulation.

***Ev-Deg-High* EVO** experiments also produced examples of functional origination and evolution from genes in the *E. coli* ORF library. One outcome was *YjiE*, which has been described as a hypochlorite-specific transcription factor in *E. coli* (*72*). Similar to WT *RPS31*, WT *YjiE* had undetectable activity, while a variant containing only one amino acid substitution (F143S, *Variant 1*) potently inhibited Ubr1-mediated degradation of N-end rule substrates (**Fig. 5B** and **Fig. S10**). However, unlike *RPS31 Variant 1*, the evolved *YjiE* variant also inhibited degradation of the GFP-PEST reporter (**Fig. 5C**), which is ubiquitinated by the SCF complex (*73*) and not Ubr1. Therefore, the evolved *YjiE Variant 1* may inhibit a component of global protein degradation or alter expression of components common between both SCF complex and Ubr1-based degradation pathways.

Perhaps the most surprising example of all ***Ev-Deg*** experiments was the fixation of lineages descended from a short *E. coli* cryptic prophage gene called *YdaG*. The evolved variant we isolated contained mutations A17T, V18M, C30Y, V38M (*Variant 1*) and substantially outperformed WT *YdaG*, which conferred no activity in selection for Ade2 stabilization (**Fig. 5A** and **Fig. S16**). This again reflects evolution’s capacity to generate gene function *de novo*. Strikingly, the evolved *YdaG* variant inhibited the Ubr1-mediated degradation of N-end rule substrates even more potently than the endogenous *ROQ1* gene, which naturally redirects Ubr1 activity, in the UbiX-Luciferase reporter assay (**Fig. 5B)**. The evolved *YdaG* variant also increased the level of a UbiY-mRuby and slightly increased the abundance of the misfolded protein degradation reporter, PHO8*-GFP (**Fig. 5C** and **Fig. S16C**), but had no detectable effect on degradation of the GFP-PEST reporter that goes through degradation by the SCF complex rather than Ubr1 (**Fig. 5C**). Therefore, the evolved *YdaG* variant likely specifically inhibits *Ubr1*.

One mechanism we considered is that the evolved *YdaG* and *YjiE* simply encode misfolded proteins that accumulate and saturate the proteasome or related machinery, reducing capacity for N-end rule substrates. However, we ruled out this possibility based on RNA-seq data, which showed no upregulation of major heat shock proteins (*74*), proteasome core subunits, or autophagy-related genes typically induced during proteotoxic stress (**Fig. S17**). Instead, RNA-seq experiments showed that strains expressing *YdaG Variant 1* and *YjiE Variant 1* exhibit a significant and specific downregulation of *PTR2* expression (**Fig. 5D** and **Fig. S18**). *PTR2* encodes a transmembrane peptide transporter that serves as a well-characterized downstream indicator of Ubr1 activity (*75*, *76*). Transcription of *PTR2* is repressed by the transcription factor Cup9. The Ubr1 E3 ligase actively targets an internal degron in Cup9 for proteasomal degradation. Hence, under normal conditions, this continuous destruction of Cup9 relieves the *PTR2* promoter from repression, resulting in enhanced *PTR2* expression. Conversely, when Ubr1 or the global protein degradation is inhibited, Cup9 rapidly accumulates in the nucleus, strongly shutting down *PTR2* expression. Therefore, the observed downregulation of *PTR2* is consistent with the stabilization of the Cup9 repressor, further supporting the hypotheses that the evolved variants of *YdaG* and *YjiE* inhibit Ubr1 and global degradation machinery, respectively. It is worth noting that while Ubr1 targets other transcription factors for degradation, those are known to be largely protected by chaperone proteins (*77*). Hence, Cup9 acts as the primary, unshielded sensor for Ubr1 activity with *PTR2* downregulation as its RNA-seq readout.

The final example from ***Ev-Deg-High EVO*** selections to extend Ade2’s half-life from the UbiR-Ade2 fusion was the emergence of a mutated chaperone gene from *E. coli*, *SecB*. The evolved *SecB* variant we isolated contained the mutations A14T, A41T, Q49L, D59N, V80I, and A145T (*Variant 1)* (**Fig. 5A** and **Fig. S16A-B**). Reporter assays on the evolved *SecB* variant’s effect on N-end rule substrates and other degradation tags did not reveal much other than a slight decrease in degradation of the GFP-PEST and PHO8*-GFP misfolding reporters. However, the effect of the evolved *SecB* variant on growth rate in **COMP**_mut/NS_ assays was high (**Fig. S16A**). The WT *SecB* also had activity (**COMP**_WT/NS_) but was much less potent compared to the evolved *SecB* variant (**COMP**_mut/WT_). The evolved *SecB* variant also slightly increased mRuby stability from the UbiY-mRuby fusion (**Fig. S16D**). Moreover, RNA-seq data revealed that the strain overexpressing *SecB Variant 1* displayed no significant upregulation of genes characteristic of a misfolded protein response (**Fig. S17I-L**), and only modest downregulation of *PTR2* compared to the control strain (**Fig. S18**), further suggesting that Ubr1 is not strongly inhibited in this case. The lack of strong activity towards alternative clients in our reporter assays might suggest that this chaperone protein primarily evolved to use the Ade2 protein as a specific client, but we do not have direct evidence for this hypothesis.

## Conclusion

In this work, we have presented an abundance of examples where new gene functions evolved from unexpected sources, including cases where the parental gene originating a novel trait had neither known function nor initial activity for the function it ultimately evolved. Our results therefore support the growing view that gene function can commonly spring from unexpected, uncharacterized, or cryptic sequences, which, although appreciated through bioinformatic and evolutionary analyses, has been difficult to experimentally witness. By developing the ORACLE platform – where large repertoires of sequences are available as starting points for rapid continuous evolution *in vivo* and where the systems level complexity of cells can provide multiple routes to selectable phenotypes – we were able to consistently capture and study the emergence and evolution of new gene function. The success of our experiments suggests that functional sequences are not as rare as evolutionary dogma has historically implied when provided the multi-gene, multi-objective, and multi-environment possibility landscapes for adaptation that define real biology.

The evolutionary outcomes we observed provide direct insights into neofunctionalization. In neofunctionalization, duplication of an existing gene affords the new copy flexibility in evolving a function distinct from its original. However, there are two categories of neofunctionalization models that differ in their details. In classical models, gene duplication occurs first, followed by mutations that sample a new function in the novel copy of the gene, which then drives its adaptive divergence (*4*). Criticisms of these models largely rely on the argument that it would be more likely for a duplicated gene to become mutationally disabled beyond repair before finding a functional mutation. However, this concern is mitigated by the fact that some of our experiments show how only a single mutation in a gene (*e.g.*, *RPS31* and *YjiE*) is needed to discover an unexpected new function that brings the gene under selection. An alternative category of neofunctionalization models suggests that genes first gain (or already possess) promiscuous activities that allow them to enter the view of selection under an environmental change that favors the promiscuous activity (*5*, *15*, *34*). Then, that activity is amplified through a beneficial duplication event (*5*, *15*), creating a new gene that adaptively diverges to transform the (weak) promiscuous activity into a strong new primary activity. The fact that we observe cases where the WT version of a gene already has a promiscuous (often unexpected) function satisfying the selections we impose, which then guides further adaptation and divergence, shows that this model of neofunctionalization may also be common. In addition to our study’s relevance to neofunctionalization, our observation that a *de novo* peptide appendage generated through a frameshift in *RTS3* originates a new metal sequestration activity lends evidence for *de novo* gene birth models where entirely non-gene sequences become new genes (*8–11*), since the *de novo* peptide appendage can be considered mostly random. Overall, our experiments support the notion that these multiple models for new gene formation are all viable, not necessarily because the merits and shortcomings of each are balanced, but because it may be fundamentally “easy” to achieve biomolecular innovations *in vivo*.

Our study is not without limitations. For example, the functions we attempted to evolve, while numerous, represent a small fraction of possible biological traits one can potentially select and did not include chemically challenging functions like enzymatic activity. Furthermore, the types of sequences we offered as substrates for evolution were restricted to extant ORFs (including dubious ORFs), and thus omitted many potential sources of gene birth, including many noncanonical ORFs, aberrantly transcribed or translated ORFs, intergenic sequences, and in the extreme, random sequence. Moreover, the scale of our study, while large (55 evolution experiments with over a dozen yielding clear examples of successful gene evolution (**Table S1**)), is still anecdotal. Addressing these limitations by broadening the range of selected functions, expanding the repertoire of evolutionary source sequences beyond extant ORFs, and increasing the cumulative number and diversity of ORACLE experiments may ultimately let us sample the true distribution of functional gene space in biologically rich *in vivo* settings. This promise, along with the utility of ORACLE in discovering the function of unknown genes and its potential use to evolve new peptides and proteins for specific applications, motivates continued ORACLE experiments that seek to understand the versatility of gene evolution and biomolecular innovation.

## Author Contributions

Conceptualization: AP, CCL; Methodology: AP, AT, CCL; Investigation: AP, AT; Visualization: AP, AT; Funding acquisition: CCL; Supervision: CCL; Writing – original draft: AP, AT, CCL; Writing – review & editing: AP, AT, CCL

## Acknowledgements

We thank Professor Wayne Patrick for his valuable contribution of the pooled ASKA library and members of the Liu group for thoughtful discussions.

## Data and Materials Availability

All data and materials generated for this study are included or available upon request to the corresponding authors. The RNA-seq data are deposited in GEO under accession number GSE335699.

## Supplementary Materials

**Materials and Methods**

**Supplementary Text 1:** ORACLE subroutines

**Supplementary Text 2:** Testing of the ORACLE experimental pipeline

**Supplementary Text 3:** Origins of libraries

**Supplementary Text 4:** Comparison of **MATE** and **PCR + INT**_p1_ for genomic background refreshing

**Supplementary Text 5:** Description of non-active gene phenomena from mating-based evolutions

**Supplementary Text 6:** Design of a good selection

**Supplementary Text 7:** Future technology developments for ORACLE

**Supplementary Text 8:** Expanding ORACLE’s scope to heterologous targets

**Fig. S1.** Optimization of emulsion PCR and demonstration of decreased size bias in **PCR**_lib_.

**Fig. S2.** Diagram of the transformation method assessment and competition assay setup.

**Fig. S3.** Visual representation of the abortive mating technique **MATE**.

**Fig. S4.** Demonstration of ability to fish out a specific combination of genes at a frequency of <1/10,000,000.

**Fig. S5.** Schematic of **PCR**_lib_ + **INT**_p1_.

**Fig. S6.** Control experiment for the isolation of a positive control gene in ORACLE.

**Fig. S7.** Full competition data from the pantothenate evolutions (***Ev-Tol-Pan***).

**Fig. S8.** Full competition data from the manganese tolerance evolutions (***Ev-Tol-Mn***).

**Fig. S9.** Full competition and reporter assay data supporting conclusions made in the main text relating to *ROQ1* and *RPS31* variants (***Ev-Deg-High***).

**Fig. S10.** Full competition and reporter assay data supporting conclusions made in the main text relating to *YjiE* and *AaT* variants (***Ev-Deg-High***).

**Fig. S11.** Full competition data from the lithium tolerance evolutions (***Ev-Tol-Li***).

**Fig. S12.** Full competition and reporter assay data from the pPHO5 evolutions (***Ev-TF***) supporting **Figure 3.**

**Fig. S13.** The evolved *YeiG* and *YdfC* upregulate the expression of many canonical Pho4-responsive genes.

**Fig. S14.** Full competition data from the copper tolerance evolutions (***Ev-Tol-Cu***).

**Fig. S15.** Full competition and reporter assay data from the medium strength degron evolutions (***Ev-Deg-Low***) supporting Figure 4.

**Fig. S16.** Full competition and reporter assay data supporting conclusions made in the main text relating to *YdaG* and *SecB* variants (***Ev-Deg-High***).

**Fig. S17.** RNA-seq data demonstrate the lack of misfolded protein response induction.

**Fig. S18.** RNA-seq data representing *PTR2* expression changes.

**Fig. S19.** Competition data from *BAX* and *AFT1* overexpression selections.

**Table S1.** Summary of all ORACLE experiments performed in this study.

**Table S2.** Genes found from successful evolutions with sequences of assessed mutants and WT.

**Table S3.** Primers used for cloning libraries and important techniques.

**Table S4.** Plasmids used for important techniques.

**Table S5.** Yeast strains generated for this study.

References (78-106)

