## Supplementary material for "Continuous evolution of gene libraries towards arbitrary functions reveals the versatility of biomolecular evolution *in vivo*": Fig. S

### Materials and Methods

**Emulsion PCR ($PCR_{lib}$).** The commercially available Micellula emulsion PCR kit (E3600-01, Roboklon, Poland) was used for preparation of emulsion PCRs. We followed the recommended ratios and volumes for each reaction. For each reaction, 220 µL emulsion component 1, 20 µL emulsion component 2, 60 µL emulsion component 3 were mixed by pipetting. When preparing multiple reactions, the volumes were scaled up and distributed in 300 µL aliquots in 1.5 mL Eppendorf tubes. The oil emulsion mix was cooled on wet ice while the aqueous PCR component of the reaction is prepared. 55 µL of a PrimeSTAR GXL DNA polymerase (TaKaRa) reaction was prepared with the modifications of addition of a final concentration 0.025 mg/mL acetylated BSA. For each library, four emulsions were prepared with a serial 1:2 dilution of template DNA to ensure that the desired occupancy of micelles was hit, starting from 10 ng of plasmid. 50 uL of the PCR mix was added to the cooled emulsion mix, and the remaining 5 µL of the PCR was saved as an open PCR control. The cooled emulsions were manually vortexed in a cold room at 4C for 5 minutes. The 350 µL reaction volume was distributed into an 8-tube PCR strip with 50 µL / well, and a well-formed emulsion makes an off-white "creamy" mix. 25x cycles of PCR were run with a $T_m$ of 60 ºC, and 2-minute extension time, alongside the open controls. Post reaction the emulsion reactions were pooled, and emulsions were broken and purified using the Supplied purification reagents and columns. 1/10 of the reaction was run on an agarose gel alongside the open control, and the emulsion reaction which yielded the lowest DNA output (lowest micelle occupancy) was selected for Gibson assembly. Primer sequences and plasmids used can be found in **Table S3, S4** respectively.

**Frozen competent yeast cell preparation**. Each desired strain was inoculated into selective media two days prior to frozen competent cell preparation (*78*). The day prior, cells were back diluted at 30 ºC with shaking at 200 rpm to be in late exponential phase the morning of. Note that for error-prone, p1-containing strains, several passages may be required for yeast to acclimate and grow well. The $OD_{600}$ of the yeast culture was measured using a spectrophotometer. Approximately 2.5 x $10^9$ cells were inoculated into 500 mL of YPD, and the culture was grown in the shaking incubator at 30 ºC for 4-6 hours until reaching a cell density of 2 x $10^7$ cells/mL. Yeast cells were harvested by centrifugation at 3,000 x g for 5 minutes and washed with 0.5 volumes of sterile water, followed by a second wash with 0.01 volumes of sterile water. The cell pellet was resuspended in 0.01 volumes of sterile-filtered frozen competent cell solution (5% glycerol, 10% DMSO, both from ThermoFisher), and 50 µL aliquots were dispensed into 1.5 mL microcentrifuge tubes. These tubes were placed in a styrofoam container and stored at -80 ºC.

**Yeast transformation**. Frozen competent cells were thawed in a 37 ºC water bath for 30 seconds and pelleted by centrifugation at 13,000 x g for 2 minutes. Frozen competent cell transformation mix (260 µL of 50% w/v PEG 3350 (ThermoFisher), 36 µL of 1 M LiAc (ThermoFisher), 50 µL of 2.0 mg/mL single-stranded carrier DNA (ThermoFisher), 14 µL of DNA plus sterile water) was added to the cell pellet, and the mixture was vortexed vigorously to resuspend the cells. The tube was then incubated in a 42 ºC water bath for 30 minutes. After incubation, the cells were pelleted by centrifugation at 13,000 x g for 30 seconds, and the supernatant was removed. Cells were either resuspended in selective media for outgrowth, or streaked on selective plates for clonal isolation. When genomic manipulation was carried out to construct a new strain, the modified locus was confirmed by PCR. Yeast strains generated in this study and their genotypes can be found in **Table S5.**

**Yeast culture and growth monitoring**. Yeast strains were grown in standard media, including yeast extract peptone dextrose (YPD) (10 g/L bacto yeast extract; 20 g/L bacto peptone; 20 g/L dextrose) or the appropriate synthetic drop-out media (yeast nitrogen base w/o amino acids (US Biological), drop-out mix synthetic minus the appropriate nutrients (US Biological), and dextrose).

$OD_{600}$ was regularly measured in a 96-well format. 100 µL of culture was measured per well using a Tecan Spark. To convert to a standard 1 cm path length, background was subtracted and the value multiplied by 10. This factor was determined empirically to match readings from a standalone spectrophotometer, corresponding roughly to 1.5 x $10^7$ cells/mL at $OD_{600}$ = 1.

**Plasmid library Gibson assembly**. DNA libraries amplified as described by **$PCR_{lib}$** were cloned into a backbone plasmid containing chloramphenicol resistance, I-Ceu-I restriction digest sites flanking the integration cassette, TP901 integration flanks, a p1 cytoplasmic promoter, and auxotrophic markers *HIS3* or *Leu2*. The open vector for each library was prepared by PCR with primers which add homology to the amplicon generated in emulsion PCR and a stop codon where appropriate, DpnI digested (NEB) and column purified. Gibson assembly (*79*) was carried out at an approximately 1:1 molar ratio of insert to backbone. Reaction was column purified and electroporated into ElectroMAX Dh5β cells, and 1/10,000 was plated for titer, and rest was inoculated into liquid selective media.

**Library p1 integration ($INT_{p1}$).** 2 µg of assembled library was digested with I-Ceu-I for 3 hours. Note that I-Ceu-I is sensitive to carryover of salt in the digest, so reactions are scaled to ensure a max of 20% of the volume of the digest was from miniprepped library DNA. Depending on the cell line being transformed, two transformations of frozen competent cells prepared as described (*80*) were sufficient to yield millions of colony forming units (CFU). Given the libraries contained around 5,000 genes this covers the libraries adequately. When transforming into cells with a lower-copy, error-prone polymerase, it was necessary to passage the cells in landing pad selective media 3–4 times before preparing competent cells, as they initially grew slowly and improved growth rate increases transformation efficiency. Post transformation, 1/5,000 of the total culture was plated on selective plates for p1 integration, and the rest (two transformations worth) was transferred to 15 mL selective culture immediately (no YPD recovery). $OD_{600}$ was measured at this point. The following day, the transformed cultures were saturated, and the libraries were banked as a starting point to be returned to. Libraries were banked in duplicate by spinning down 12 mL, and resuspending in 2 mL of 10% DMSO 5% glycerol and freezing in an isopropanol freezing chamber. Cells were passaged three times prior to selection as it takes approximately this many passages to stabilize expression from the integrated p1 cassettes. A GFP-only control cloned into the same backbone was included alongside the library integrations to monitor expression and as an evolution control.

**Yeast abortive mating (MATE).** One day prior to abortive mating, p1 donor library containing were passaged into media only selecting for the linear plasmid (-H or -L). p1 receiver cells which contain only genomic modifications were passaged to YPD to a final $OD_{600}$ of 0.025. On the day of mating, the $OD_{600}$ of both the p1 donor and receiver cells was measured, and 1 mL of $OD_{600}$ = 1, p1 receiver, and 0.5 mL of $OD_{600}$ = 1 p1 donor are mixed, spun down, and resuspended in 25 µL of 0.9% NaCl. This mix of cells was spotted onto a YPD plate and allowed to dry before the plate is flipped and incubated at 30 °C for 6 h. At this scale, the yield of desired cells (correct haploid with p1 transfer) can be expected to be around 500,000 CFU. The mating can be scaled up or down as desired. For abortive mating in a 96-well plate, we scaled down 5-fold and spotted 4 columns of 8 rows using a multichannel pipette to fit 32 mating reactions on a single YPD plate. After 6 h the mated cells were scraped into selective media. In our setup, for one direction of mating, this media was composed of -HR + Nourseothricin + Canavanine (no ammonium sulfate). For mating the other direction, this media was composed of -HUK + Thialysine. The following day the second layer of selection was added (-HR+Nat+Can+5FOA or -HUK+Thia+5-FC). Including 5-FOA immediately after mating reduced the CFU by 10-fold, as even desired still contain residual Ura3 in the cytoplasm, and vice-versa with 5-FC. Canavanine and thialysine are not problematic for immediate selection as they are membrane bound transporters, and membranes are resynthesized with cell budding. After mating the cells were passaged in full counter selection 2x, then once in -H or -HU, then into selective media.

**Antibiotic concentrations and media conditions for counterselection.** Thialysine was used at a final concentration of 50 µg/mL in media lacking lysine (-K). Canavanine was used at 50 µg/mL in media lacking arginine (-R). 5-FOA was used at 500 µg/mL in SC media containing uracil. 5-FC was used at a concentration of 25 µg/mL in liquid SC.

**Library evolution with retransformation (PCR + $INT_{p1}$).** For the configuration of ORACLE utilizing the retransformation, frozen competent cells were prepared as described but banked as 12.5 µL aliquots rather than 50 µL aliquots. This was done to make the transformation compatible with recovery in a 24 well block. After a round of evolution (first round starting from the full library transformation), DNA was prepared by: measuring $OD_{600}$ of each well and pipetting volume to reach an equivalent of 6 x $10^6$ cells (1 mL of $OD_{600}$ = 0.4) into an Eppendorf tube. All samples were spun down at >17,000 rpm for 15 seconds, media was aspirated completely, washed with 1 mL of dd$H_2O$, spun down again, and dd$H_2O$ was aspirated completely. This was done to ensure consistency across wells as media carryover can impact PCR efficiency. Pellets were resuspended in 100 µL of 5% Chelex, and glass beads were added to approximately ½ the volume. Tubes were vortexed for 6 minutes on a foam tube holder, and then boiled for 6 minutes on a heat block set to 105 °C. All tubes were spun down at max speed for 10 seconds, and 40 µL of DNA solution was transferred to a 96 well for storage. PCRs were conducted with Platinum Superfi 2x PCR mix (Thermo #12369010).

For PCR, a master mix of 10 µL 2x PCR mix, 6.75 µL of dd$H_2O$, and 0.125 µL 100 µM primers, (scaled up to reaction number) was prepared and distributed to the appropriate number of PCR tubes (16 uL/rxn). 3 µL of GC prepped DNA was distributed to prepared master mix. 17x cycles of PCR with a 4-minute extension time (theoretically sufficient for 16kb) were run. 2 µL of PCR was used to assess PCR on an agarose gel. 13 µL of unpurified PCR mix was used as DNA template for transformation at 0.25x scale the regular transformation. With well prepared competent cells, this yields about 100k transformants / transformation. Post transformation the cells were recovered in 2 mL -H overnight and passaged twice prior to selection. The number of passages in selective evolution media should be determined according to how quickly a genomic adaptation can take over the culture.

**Evolution conditions (EVO).** For all evolution conditions, the libraries were initially passaged to a starting $OD_{600}$ of 0.1 in 2 mL (approximately 3 x $10^6$ cells), and as the active sequences enriched, the amount of cells passaged was decreased, first to 0.05, and then to 0.025 $OD_{600}$. Cells were passaged every day. Reducing the cells passaged allowed for faster enrichment at the cost of diversity.

- **Manganese:** Cells were passaged in 25 mM $MnCl_2$ media in -H. After the first round of evolution the concentration of $MnCl_2$ was increased to 30 mM, and then 35 mM.
- **Pantothenate dropout:** Media lacking pantothenate was prepared with YNB lacking calcium pantothenate (Formedium CYN3401).
- **Copper:** Media containing 2.5 mM Copper Sulfate, was gradually increased to 2.75 mM, then 3.0 mM.
- **Lithium:** Media containing 250 mM Lithium Chloride was used.
- **Adenine dropout:** Transformations and regular culture was conducted in -H++A, media supplemented with 80 mg/L adenine. For selective conditions, media lacking histidine and adenine was prepared, and gradual dropout of adenine was carried out, first to 2.5 mg/L, then 1.25 mg/L, then 0 mg/L of supplemental adenine.

**Fluorescent competition ($COMP_{X/Y}$).** Competition strains which contained a Zeocin TP901 genomic landing pad, GFP or mRuby fluorescence cassette, and TP901 2μ plasmid were prepared as frozen competent cells. DNA was prepared from each evolution campaign by amplifying genes with primers that add 45 bp of homology to integration flanks. Integration flanks were prepared by PCRing a plasmid containing a TP901 site-promoter-terminator-*HIS3*-TP901 site. Integration fragments were specific to the library as the cloning scar leftover from the original library differs based on the vector the library was cloned from. PCR of genes was done as for preparation of the **PCR+INT**$_{p1}$ based evolution process. 5 μL of each flank and 5 μL of the variable genes were mixed and transformed in a ½ scale frozen comp cell transformation (**INT**$_{genome}$). For example, the GFP strains were transformed with genes from evolution, and the mRuby strains were transformed with a nonsense control.

After 2 days recovery, cells are passaged once with a 1:50 dilution. The following day the cells of interest were mixed with the control (nonsense integration or WT) 1:1 by volume. The $OD_{600}$ of the mixture was noted to use for calculation of the starting $OD_{600}$ in competition. The mixture of cells was inoculated at a 1:100 dilution (or depending on media condition) into selective media, and the ratio at day 0 was measured. The following day, $OD_{600}$ was remeasured and flow cytometry was used to measure the cell ratio again, and the cells were again passaged noting the $OD_{600}$ to monitor population expansion, and so on. A nonsense vs. nonsense control was run alongside the competition as eventually the competition becomes unstable due to genomic adaptation unrelated to the transformed gene. 20,000 cells were measured per well by flow for GFP/mRuby ratio in competition.

**Calculation of relative growth rate and enrichment values from competition.** We report relative growth rate values from the competition experiments unless otherwise indicated. These can be simply calculated by using the population expansion of the two species in the same well. We measure the fractions of cells that were green ($GFP^+$) and red ($mRuby^+$) in the first timepoint, the starting $OD_{600}$, and final ratio of green/red cells in the end timepoint, and $OD_{600}$. As the $OD_{600}$ is directly proportional to the number of cells in the culture, we calculate:

$$Generations(Green) = \log_2 \frac{(\%Green_f * OD_f)}{(\%Green_i * OD_i)}$$

The same calculation is done for the red strain within the same well. This is converted to growth rate by:

$$Relative\ growth\ rate(Green/Red) = \frac{Generations(Green)}{Generations(Red)}$$

In some of the supplementary figures we report the enrichment calculated over different passages. In a scenario where the evolved GOI and nonsense control are being expressed in the green and red strain respectively, we calculate:

$$log_2\,(Enrichment) \; = \; log_2 \frac{\left(\frac{Fraction\ Green_{T_x}}{Fraction\ \ Red_{T_x}}\right)}{\left(\frac{Fraction\ Green_{T_0}}{Fraction\ \ Red_{T_0}}\right)}$$

**ICP-MS for determination of intracellular copper concentration.** Yeast strains expressing wild type or the evolved versions of the *RTS3* gene under the RPL18b promoter encoded at the HO locus were built. These strains were passaged twice in media containing 2.5 mM copper sulfate. 25 ml of saturated yeast culture was collected by centrifugation at 3000 x g for 10 minutes at 4 °C. The harvested yeast pellet was washed twice with 0.9% sodium chloride containing 5 mM EDTA. Then, the cells were washed 4 times with 0.9% sodium chloride to remove EDTA and the remaining extracellular copper. After each wash, the supernatant was collected using a micropipette without disturbing the pellet. The harvested cells were resuspended with 250 ul of milliQ water and the optical density was measured using a spectrophotometer. The remaining sample was autoclaved for 20 minutes at 121 degrees and sent to Solvias, Kaiseraugst, Switzerland for ICP-MS. The samples were decomposed by addition of oxidizing acids and heating in a closed-vessel, microwave-heated, high-pressure autoclave. The concentration of Cu was determined using an Agilent 7900 instrument. The quantitation limit/ reporting limit for the measurement was 0.1 mg/kg.

**96 well frozen competent cell preparation and transformation**. Frozen competent cells were prepared as described above with a couple key changes to facilitate easier transformation with no centrifugation. The same outgrowth conditions were used, but one extra water wash step was conducted. Cells were frozen at the same concentration of competent cell mix, but 4 µL was aliquoted per well of a 96 well plate and frozen in a styrofoam box. On the day of transformation, the ssDNA + PEG + LiAc master mix was prepared to 45 µL / well containing 32.5 µL PEG, 4.5 µL LiAc, 1.25 µL ssDNA, 2.25 µL $H_2O$, 1.5 µL of the PCR gene amplicon, and 1.5 µL of each flank. For transformation in 96 well format, plates were thawed by placing at 37 °C for 1 minute, and foil was removed. Thawed cells in DMSO / Glycerol mix were resuspended with 45 µL of transformation mix by pipetting (note, these were not spun down and frozen cell mix was left in the transformation) and then incubated at 42 °C for 30 minutes in a thermocycler. Post heat shock, the mixture of cells + PEG was diluted to 1 mL in SC media + selection conditions by pipetting (again note, the PEG + DNA mixture was not removed from the outgrowth). The 1 mL was split into two wells for bioreplicates as the transformed cells would be separate transformation events. These were grown for two days and 1/25 of desired wells were plated for titers. Cells were saturated after two days and generally, 1-5,000 transformants per well was obtained with the genomic TP901 landing pad method.

**Luciferase assay**. Nanoluciferase containing reporter strains transformed with the 96 well method were outgrown two days to saturation, passaged twice more at 1:500 for 24 h each passage. On the morning of luciferase measurements, cells were passaged 1:20 in -H media. After 5 hours of outgrowth $OD_{600}$ of wells was measured. $OD_{600}$ of all wells was within a 10% range indicating all were in exponential growth. Cells were diluted 1:3 in 0.9% NaCl to a final $OD_{600}$ of approximately 0.5-0.6. 8 µL of diluted cells was transferred to a white, opaque 384 well plate, with no two wells adjacent to another occupied well to minimize cross talk during light detection. Samples were mixed with Promega nano luciferase kit (N1110) 1:1. Luminescence was read on Biotek Synergy H1 with an integration time of 1 second, gain 135. Nonsense control was included as a sample in plates to ensure normalization was accurate. Luminescence was first normalized by $OD_{600}$, and luminescence was reported as $log_2$ (RLU sample / RLU control) for the appropriate cell line.

**GFP and mRuby degradation assays**. Fluorescent reporter containing strains transformed with the 96 well method were grown two days to saturation, passaged twice more at 1:500 dilutions for 24 h each passage. Cells in late log phase were diluted in 0.9% NaCl with propidium iodide and gated for viability, FSC/SSC, and FSC-A/FSC-H for single cells. 20k cells per well were probed and $\log_2$ (MFI Geomean sample / MFI Geomean control) was reported in these assays.

**Yeast growth curves and max growth rate calculation**. To measure the max growth rate, yeast were grown to saturation, then passaged with a 1:500 ratio twice (once per day). On the morning of the growth curve measurement, yeast were back diluted 1:10 and grown for 4 hours to ensure that all cells were in exponential growth prior to inoculation. Cells in exponential growth were inoculated 1:100 (1 µL into 100 µL of -HA) of a clear bottom 96 well plate. Samples were measured in biological duplicate. The plate was covered with a breathable film and transferred to a Biotek plate reader shaking at 500 rpm, 30 °C, with $OD_{600}$ measured every 15 minutes. Growth curves were complete within 24 hours of growth. To calculate the growth rate, the natural log of the $OD_{600}$ values was calculated and the slope of the linear range was fit using Graphpad software.

***E. coli* growth curves and max growth rate calculation**. To measure the max growth rate, *E. coli* containing IPTG-inducible plasmids were grown to saturation, then back diluted 1:20 in 2YT+IPTG and grown for 2 hours to ensure that all cells were in exponential growth and expressing the plasmid prior to inoculation in the copper media. Cells in exponential growth were inoculated 1:100 (1 µL into 100 µL of 2YT + 4 mM copper chloride) into a clear bottom 96 well plate. The plate was covered with a breathable film and transferred to a Biotek plate reader shaking at 500 rpm, 37 °C, with $OD_{600}$ measured every 15 minutes. The growth curves were complete within 24 hours of growth. To calculate the growth rate, the natural log of the $OD_{600}$ values was calculated, and the slope of the linear range was fit using Graphpad software.

**Fluorescence-Activated Cell Sorting for overexpression evolution**. A base strain expressing *GOI*-2A-GFP (where GOI is *BAX*, *AFT1*, *HSF1* or *PPZ1*) from an inducible GAL1 promoter and a TP901 p1 landing pad was constructed. The yeast and *E. coli* ORF plasmid libraries were digested and integrated as discussed above. Library cells were passaged once to 0.2% glucose, 2% raffinose synthetic dropout media and then induced in 2% galactose media lacking glucose. All samples were normalized to $OD_{600}$= 0.5 for all the passages. 2-3 days post-induction in galactose media, cells were live-dead stained using the TO-PRO-3 Stain (Thermo-fisher scientific) and sorted for 100 thousand $GFP^+$ cells. After each two rounds of induction and sorting, the p1 library was PCRed and re-integrated to fresh strains as mentioned above. This cycle was repeated for 4 sorts in total.

**RNA-seq.** Yeast strains encoding evolved genes or a nonsense gene were passaged twice in nonselective (adenine-rich) media. Cultures were then grown overnight in either nonselective (80 mg/L adenine) or selective (low 2.5 mg/L adenine) media, testing two to three biological replicates per condition. Cells were harvested at mid-log phase ($OD_{600}$ = 0.5 - 1) from 700 ul of culture. Cells were digested using zymolyase, and total RNA was extracted using the RNeasy kit (Qiagen).

Purified RNA was submitted to Plasmidsaurus for library preparation, Illumina sequencing, and downstream bioinformatics analysis. Libraries were prepared and sequenced using a 3' end-counting approach. The sequencing depth per sample averaged around 20 million raw reads. Raw reads were demultiplexed and quality-filtered using fastp (*81*) with a minimum Phred quality score of 15, and a minimum length requirement of 50 bp. Quality-filtered reads were aligned to the yeast reference genome using STAR aligner (*82*) followed by coordinate sorting using samtools. Gene-level expression was quantified with featureCounts (*83*), and differential gene expression analysis was performed using the package edgeR (*84*), based on the negative binomial distribution.

### Supplementary text

### Supplementary text 1: ORACLE subroutines

Each ORACLE experiment started with $\mathbf{PCR}_{lib}$ where we amplified complex gene libraries from various plasmid gene library collections and subcloned those PCR products into OrthoRep integration plasmids (*80*). These integration plasmids were then used to integrate their contained gene libraries onto OrthoRep's hypermutating p1 in *S. cerevisiae* stains (INTEGRATION$_{p1}$ or $\mathbf{INT}_{p1}$). This was done either directly into the selection strain for evolving the gene library or into a general donor strain that was abortively mated with a selection strain to transfer the cytoplasmic p1s *en masse* (**MATE**). Once a gene library was in the selection strain, continuous growth-based evolution experiments were carried out (**EVO**). Throughout **EVO** from time to time, the p1 plasmids comprising the population of evolving genes were amplified (**PCR**) and transferred into fresh selection strains by $\mathbf{INT}_{p1}$ or directly transferred by **MATE** to prevent the accumulation of any genomic mutations that could confound gene adaptation.

#### $\mathbf{PCR}_{lib}$

PCR required amplifying diverse multi-sized libraries of genes, which risked introducing size biases and chimeric recombinants among genes. We therefore adopted emulsion PCR to minimize these risks, optimizing the process by titrating BSA to balance emulsion stability against PCR inhibition (**Fig. S1A**), titrating DNA to verify an approximate micelle number, and ensuring emulsion integrity by differential amplification of short and long amplicons (*85*) (**Fig. S1B**). During gene library preparation where gene libraries were amplified from plasmid library sources (*41–43*), DNA titration was used to find a concentration where template-containing droplets were abundant and where most template-containing droplets contained a single template plasmid, reducing size-based PCR biases relative to pooled PCR (**Fig. S1C**). After PCR of gene libraries, we used Gibson assembly (*79*) to subclone into a p1 integration vector redesigned with I-CeuI sites (to avoid common restriction sites) for linearization and exposure of TP901 attachment sites for integration onto p1 (*80*). In the current work, three DNA libraries were each subcloned into two p1 integration vectors, one of which encodes *HIS3* and the other of which encodes *LEU2* for selection of p1's after linearization and integration, for a total of six p1 integration vector libraries. The three distinct libraries represented are 1) a yeast ORF library (*41*), 2) an *E. coli* ORF library (*42*), 3) an *E. coli* ORF library where each member is fused to a GFP tag (*42*)

#### $\mathbf{INT}_{p1}$

We previously reported a high-efficiency method for integrating genes onto p1 landing pads (*80*). The method consists of a p1 landing pad with attachment sites for TP901, donor DNA that is flanked by corresponding attachment sites (*i.e.*, linearized integration vectors), and the transient introduction of TP901, which carries out the high-efficiency recombination of donor DNA onto p1. The product of integration was selected for using markers contained in the donor DNA, specifically *HIS3* or *LEU2*. The high transformation efficiency of TP901-mediated integration using fast and routine protocols ensured that the genetic diversity of libraries and throughout evolution experiments was almost certainly oversampled during $\mathbf{INT}_{p1}$ steps.

#### $\mathbf{INT}_{genome}$

Similar to $\mathbf{INT}_{p1}$, routine integration into a genomic locus was made possible using TP901. Additionally, because of yeast's efficient nuclear recombination mechanisms, overlapping fragments can be transformed, where they will assemble into the genomic locus *in vivo*. To enable rapid characterization of evolved gene populations at a non-mutating genomic locus at scale, we leveraged both TP901 and yeast nuclear recombination (**Fig. S2A**). For every evolved gene population we wished to characterize, we used **PCR** for amplification from p1, attaching ~60 bp overlaps to generate common flanks in the process. The 5' gene overlap was homologous to the 3' overlap of a synthetic DNA fragment that encoded a constitutively active nuclear promoter (usually pRPL18B (*51*)). The 3' gene overlap was homologous to the 5' overlap of a synthetic DNA fragment that encoded a selection marker (*e.g.*, *HIS3*). The 5' end of the nuclear promoter fragment contained the 5' TP901 attachment site, and the 3' flank of the selection marker fragment contained the 3' TP901 attachment site. These three fragments were transformed into strains engineered with a genomic landing pad locus with corresponding TP901 attachment sites and containing a plasmid expressing TP901 recombinase (*86*). Fragments reliably recombined and integrated at the TP901 attachment sites when transformed, resulting in ~100,000 successful transformants per routine experiment where each transformant is a single gene mutant encoded at a copy number of one.

**COMP**$_{X/Y}$ (where X and Y are what are being competed against each other)

Given the large scale at which we ran ORACLE evolution experiments, we needed streamlined characterization pipelines to accurately measure the relative performance of evolving populations against appropriate controls. To satisfy this need, we developed a high-resolution competition assay. In this assay, variable gene fragments from p1 are amplified by **PCR** and integrated by **INT**$_{genome}$ into two otherwise isogenic yeast selection strains marked with two different fluorescent markers, GFP and mRuby (**Fig. S2A**). One strain (*e.g.*, GFP-tagged strain) then receives evolved gene populations or an evolved gene mutant, while the other (*e.g.*, mRuby-tagged strain) is transformed with a control fragment (*e.g.*, a short nonsense sequence or the WT ancestor of an evolved gene). Mixed cultures (1:1 ratio) are then grown under selection for 24 h per passage (**Fig. S2A-B**). The initial ratio and final ratio of GFP versus mRuby cells are determined by flow cytometry (representative flow plots in **Fig. S2C**) and population sizes are measured by optical density ($OD_{600}$). Given time, change in $OD_{600}$, and GFP versus mRuby ratios, we calculate an apparent growth rate constant for each strain (see **Materials and Methods**). We call this an apparent growth rate constant, because growth may not be continuously exponential over 24 h. Since these growth rate measurements are done in pairwise competition against a control, comparison of an evolved gene population or evolved gene mutant against the control is expected to be maximally reliable. **INT**$_{genome}$ was confirmed to generate pure populations of yeast overexpressing the gene of interest (**Fig. S2D**) ensuring that competition is among pure populations. Additionally, we often also compete strains containing nonsense fragments against each other, *i.e.* **COMP**$_{X/Y}$. This is to ensure that the two differently marked fluorescent strains have the same growth rate (*i.e.*, GFP or mRuby expression affects both strains equivalently).

**MATE**

We developed an abortive mating-based system to efficiently transfer p1 plasmids into new hosts, for example, to move gene libraries into selection strains or refresh p1's host genome background during evolution experiments. OrthoRep's p1 plasmid is maintained cytoplasmically. Abortive mating allows yeast cytoplasms to mix while avoiding nuclear fusion due to a defective *KAR1* allele (*87–89*) (**Fig. S3**). Abortive mating is expected to produce three cell types containing the p1 plasmids of the donor: the original donor haploid, spurious diploids, and the recipient haploid, approximately in a 2:1:2 ratio (*89*). We selected for the desired recipient haploid population containing p1 plasmid using a combination of positive selections on auxotrophic and/or antibiotic resistance markers along with layered counterselections (*e.g.*, thialysine, canavanine, 5-FC, 5-FOA, 5-FAA) to eliminate donor cells and diploid "cheaters" (*90–94*), as diagramed in **Fig. S3** and detailed in **Materials and Methods**. Note that the donor and recipient strains are opposite mating types such that each mating round changes the mating type of the host cell.

**MATE** may also be useful for evolution experiments where multiple p1 species encoding different genes are maintained by distinct markers for gene coevolution. This is because **MATE** allows a mixture of p1s from one cell to transfer into another cell without changing the composition of the mixture. Although we do not carry out coevolution of multiple p1-encoded genes in this work, we have validated that cells can stably maintain and express multiple p1 plasmids selected using different markers where sequential **MATE** is used to generate populations containing p1 mixtures in each cell at high enough efficiency where all pairwise combinations of ORF libraries can be constructed (**Fig. S4**). For detection of extremely rare events (*e.g.*, a specific pair of genes) it was necessary to apply a technique called MERGE (*95*), wherein the p1 destination strain contains a Cas9 plasmid and guide targeting the locus to be converted to match the destination strain (**Fig. S4**).

**PCR** + **INT**$_{p1}$

An alternative to **MATE** as a means of refreshing p1's host genome background during **EVO** is to rely on **PCR** + **INT**$_{p1}$ (**Fig. S5A**). By **PCR** amplifying p1's evolving gene locus with the addition of flanks that include attachment sites for TP901-mediated integration, it was straightforward to then use **INT**$_{p1}$ to directly reintegrate the amplified gene population into fresh selection strains containing 1) a p1 landing pad with TP901 attachment sites and 2) a transiently maintained plasmid expressing TP901. PCR bias (especially against longer genes) and formation of recombinants were minimized by reducing PCR cycle numbers (*85*). An advantage of **PCR** + **INT**$_{p1}$ is that nonfunctional p1 genes that can be generated from and hitchhike with evolving functional genes in the same cell (*96*) – p1 is multicopy – may be rapidly purged after **INT**$_{p1}$, as only one or a few copies of donor DNA are integrated into each cell during TP901-assisted integration onto p1 (*80*). The use of **PCR** + **INT**$_{p1}$ from time to time during **EVO** may therefore not only afford a procedurally straightforward way to refresh p1's host genome background but may also enable faster selective sweeps by beneficial mutants.

Optimizations that streamlined **PCR** + **INT**$_{p1}$ so that it could be carried out rapidly at scale included using GC-prep isolation to extract p1 populations (*97*), quarter-scale transformations (~100,000 CFU per transformation), and frozen competent cell preparations (*78*). Together, these optimizations allowed for up to 96 **EVO** campaigns that included **PCR** + **INT**$_{p1}$ steps to be carried out concurrently by one researcher.

### Supplementary text 2: Testing of the ORACLE experimental pipeline

To test whether the setup steps leading to **EVO** in the ORACLE experimental pipeline were reliable, we first carried out a mock ORACLE experiment where we simply sought to enrich a known selectable gene from a p1-encoded yeast ORF library encoding ~5,200 yeast ORFs (*41*) (**Fig. S6**). **INT**$_{p1}$ was used to install the gene library into an OrthoRep strain that uses a WT orthogonal DNAP (WT TP-DNAP1) to replicate p1. The resulting population was then transferred into a selection OrthoRep strain using **MATE**. This strain encoding an error-prone DNAP (Trixy) (*40*) for p1 replication had its *ARO7* gene deleted, rendering it a phenylalanine auxotroph (*98*) that could be used to guide the evolution of the hypermutating p1-encoded genes to complement the auxotrophy. Since the *ARO7* gene is in the yeast ORF library installed onto p1, we expected that **EVO** in the absence of phenylalanine would enrich p1-*ARO7* library members. We found this to quickly occur simply after plating cells in media lacking phenylalanine where almost all surviving colonies contained p1-*ARO7*. This proved the integrity of the **PCR**$_{lib}$, **INT**$_{p1}$, and **MATE** subroutines and demonstrated that a known gene function could be selected by **EVO**.

### Supplementary Text 3: Origins of libraries

The yeast ORF library was sourced as an arrayed library from Horizon Bio (#YSC3868), which contains all known and dubious yeast ORFs. This library was previously cloned in an arrayed format and is supplied in *E. coli* using a 96-well plate (*41*). To convert it into a pooled format, each well was grown separately, then pooled, and DNA was prepared from the pooled cultures. Several genes—*ADE2, TRP5, URA3, HIS3, KAR1, LYS2, CAN1, LEU2, LYP1*, and *MET15*—were intentionally omitted to facilitate selections. The PCR primers include overhangs for Gibson assembly, and the reverse primer introduces a stop codon three amino acids downstream of the ORF to remove a tag from the original library. For competition experiments, individual wells containing the gene of interest were regrown to serve as PCR templates. Cloning and competition primers are listed in **Table S3.**

Next was an *E. coli* ORF library, previously constructed in an arrayed format (*42*). The pooled plasmid library was generously gifted to us by Professor Wayne Patrick, as he had previously constructed the pooled library (*43*). This library contained the *E. coli* ORFs tagged at the C-terminus with GFP, and a Hisx6 tag at the N-terminus. This was cloned in two formats, one omitting the GFP by amplifying from the linker connecting the gene to the GFP and introducing a new stop codon, and one including the GFP. Note that the mutations reported in the paper are in frame with the Hisx6 tag. To convert the mutations to their untagged gene, subtract 16 from the position of the residue listed. The former was cloned for concerns that the connected GFP may reduce the activity of some proteins, and the latter as we guessed that the GFP may help some *E. coli* genes to be soluble in yeast. Note that in the competition experiments to obtain the WT gene for comparison, we had to order primers for each gene specifically. For these, PCR was carried out on the original ASKA-GFP library with gene specific primers and the band of the correct size was gel extracted and re-amplified with primers, which add handles such that they can be amplified with the same primers as are used for the amplification of *E. coli* genes. For the WT version of GFP tagged genes, the WT genes were amplified with gene specific primers, and then PCR assembled with GFP to transform in the same format as the evolved genes. Having only one specific primer was problematic for amplification and would result in a smear. Cloning and competition primers can be found in **Table S3**.

### Supplementary Text 4: Comparison of MATE and PCR + $\mathbf{INT_{p1}}$ for genomic background refreshing

In early ORACLE experiments, **MATE** was used to refresh strain's genomic background (**Table S1**). In these experiments, we performed evolution in alternating rounds: an initial round conducted in strains expressing each of the error-prone DNAPs, followed by a round in strain expressing wild-type DNAP to replicate p1 and impose purifying selection, and then an additional round in error-prone DNAP strains to further diversify the ORF libraries.

In subsequent ORACLE experiments, the same pipeline was used, consisting of $\mathbf{PCR_{lib}}$, $\mathbf{INT_{p1}}$, and **EVO** but with the modification that instead of **MATE** to transfer p1 into fresh selection strain backgrounds at the end of each round we used **PCR** + $\mathbf{INT_{p1}}$ (see **Table S1** for details). This effectively accomplishes the same operation as **MATE** and is a similarly streamlined process (see **Supplementary Text 1**), with two advantages. First, we suspected that **MATE** strains could create minor selection pressures unrelated to the selection conditions for **EVO** (**Supplementary Text 5**), which we wished to avoid all else equal. Second, we note that p1 is a multicopy plasmid so each cell during **EVO** could contain a mixture of gene mutants. We suspected that **PCR** + $\mathbf{INT_{p1}}$, which results in the installation of only one or a few copies of evolving genes into p1, would reduce the intracellular competition for fixation of gene variants encoded on different p1s and afford faster selective sweeps of superior genotypes (*96*). It's worth noting that in these subsequent experiments, **EVO** was carried out exclusively in error-prone DNAP strains, without an intervening wild-type DNAP round.

### Supplementary Text 5: Description of non-active gene phenomena from mating-based evolutions

While the **MATE** procedure was used in numerous successful ORACLE experiments, there were often genes which appeared to be adaptive, based on the growth rate of yeast containing the p1 plasmid, but did not confer a fitness advantage when tested in **COMP** experiments where they were encoded on the genome. Notably, the same gene, *SIT4*, was recurrently the fixed gene in several evolution campaigns with different selection pressures (pPHO5-*ADE2*, manganese, pPOP6-UbiY-*ADE2*). This occurred despite confirmation of the genomic *ADE2* replacement cassette post-evolution, indicating that the bypass mechanism was not due to the preservation of 'cheater' diploids. Furthermore, the "evolved" *SIT4* variants often contained frameshift mutations. The identification of *SIT4* on multiple occasions may offer a clue for these evolution experiments that did not confirm adaptation in **COMP** assays. Specifically, in the evolution strain, we knocked out the gene *SSD1*, as this has been shown to be responsible for autodiploidization, which is a common genomic mechanism of adaptation in adaptive laboratory evolution experiments. The exact role of *SSD1* is unclear but it is named for "suppressor of *SIT4* deletion." It is possible that *SIT4* was compensating for disrupted TOR signaling or cell cycle maintenance. However, these observations imply that additional, unintentional selection pressures were at work during library evolution. Conducting evolutions in cells with intact *SSD1* will likely mitigate this issue. For the purposes of this paper, rather than optimize in this mating-specific setting, we developed the **PCR** + **INT**$_{p1}$ subroutine and used cells with intact *SSD1* for all corresponding experiments.

### Supplementary Text 6: Design of a good selection

For researchers interested in applying ORACLE to uncover novel biology, we offer the following guidelines for designing an effective evolution campaign. Three main criteria should be considered:

First, the stability of the selection. If an adaptive evolution experiment can easily improve the growth rate of yeast through genome adaptations within a small number of passages (2-5 passages), that is likely to happen during the evolution campaign before genome refresh steps, and could lead to the hitchhiking-based fixation of a random gene from the ORACLE libraries, rending gene evolution moot. The stability of selection can be experimentally verified with a mock competition experiment, competing otherwise isogenic GFP+ and mRuby+ cells, and measuring the growth rate and ratio over several passages, in replicate. A good selection will have only a small change in cell ratio. Easily cheated selections will enrich the first clone which stochastically finds the genomic phenotype and along with it, an irrelevant p1.

Second, the impact of the selective condition on growth rate should be large and highly improvable. The reader may notice that we had most success using *ADE2* replacement in the main text. In these contexts, cells grew very slowly without supplemental adenine, but resolving the selection enables a fast growth rate. The pressure is also externally controllable by the addition of differing concentrations of supplemental adenine. Some selections, where global pressure is being applied (*e.g.*, ethanol tolerance) may suffer from the problem that only small increases in fitness are available (*e.g.*, yeast already grow well in moderate concentrations of ethanol). OrthoRep works best in settings where discovering and enriching an evolved variant leads to a large fitness increase, as one might expect.

Third, the impact of the selection condition on expression of p1 encoded genes should be measured. We typically include a p1-GFP-*HIS3* control carried alongside our **EVO** experiments, which enables a facile way to monitor general p1 copy number and/or expression changes due to the selection condition. As p1 relies on unique machinery for transcription and translation, some selection conditions may interfere with expression of genes. This may be problematic, as we speculate that those conditions are likely also interfering with expression of the p1 encoded auxotrophic marker, and thus some pressure will be to improve expression of p1 encoded genes through pathways that are not related to a gene library member adapting to the desired environmental pressure. While some p1 expression adaptation is not concerning, the detection of p1-GFP-*HIS3* expression change raises our level of suspicion in gauging the success of **EVO** even if growth rates increase during **EVO**.

### Supplementary Text 7: Future technology developments for ORACLE

There are several obvious technological developments that ORACLE will undergo. First, the use of standard **PCR** + $\textbf{INT}_{p1}$ during **EVO** is suboptimal, as standard PCR biases evolutionary outcomes towards shorter genes. $\textbf{PCR}_{lib}$ uses emulsion PCR to remove standard PCR biases but emulsion PCR is tedious so we only used it during the initial construction of libraries. Developing a facile method of emulsion PCR to amplify the p1 libraries during **EVO** will be worthwhile.

Next, the capacity to evolve multiple genes within the same cell, which requires **MATE** to refresh the genomic background during **EVO** while keeping coevolving genes on different p1s together in the same cell, has not been explored in this work. In its current form, **MATE** suffered from an unfortunate artifact (see **Supplementary Text 5**). This issue may be easily resolved by genetic manipulation of the evolution strain but could also be dealt with by designing selections which don't require refreshing the genome. An upside of the **MATE** approach is the ability to easily generate libraries of unprecedented size, where tens of millions of gene combinations are subjected to hypermutation with the upper library size limited only by scale of culture. The ability to maintain multiple p1's has been tested up to three p1's per cell.

Finally, the usable libraries are not limited to yeast and *E. coli* ORFs even though these are the ones that yielded all the successes in our current work. Access to new ORF libraries and development of methods to amplify libraries from cDNA will productively expand the scope of ORACLE.

### Supplementary Text 8: Expanding ORACLE's scope to heterologous targets

We were interested in exploring whether we could evolve regulators of therapeutically relevant human signaling proteins, which should be broadly tractable with ORACLE, since human regulatory proteins often maintain the same function when transferred into yeast (*99*) and yeast models have been used to study human disease (*100*, *101*). As demonstration, we set out to evolve genes to inhibit the human protein, *BAX*, a pore-forming pro-apoptotic protein of the *Bcl-2* protein family that plays a crucial part in the mitochondrial pathway of apoptosis. While reduction of *BAX* activity has been implicated in certain cancers and neurodegenerative diseases (*102*), *BAX* depletion can also be protective in diseases of uncontrolled cell death (*103*–*105*). It is known that in *S. cerevisiae*, *BAX* expression induces cell death by cytochrome c release as it does in mammalian cells (*100*, *101*). This enables a growth-based selection for *BAX* inhibition.

In an ORACLE strain, we genomically encoded *BAX* under the inducible *GAL1* promoter and coupled the expression of *BAX* to a sortable marker, *GFP*, using a 2A peptide fusion. This allowed for the selection of genes that inhibited *BAX* activity under galactose induction. Periodic rounds of fluorescence activated cell sorting (FACS) for GFP expression prevented the fixation of simple genomic mutations that disabled

*BAX* expression. We carried out **EVO** from the yeast ORF library in an experiment that included four cycles of *BAX* induction, growth, and FACS. This experiment resulted in the fixation of an evolved *NNT1* (*Variant 1*) with mutations D19N, K49R, G60S, G90D, N135D, and A219T. *NNT1* is a protein methyltransferase known to methylate elongation factor 1$\alpha$. The evolved *NNT1 Variant 1* promoted survival in the presence of *BAX* induction while the WT *NNT1* did not provide any protective effect and was indistinguishable from the nonsense control (**Fig. S19A**). While we did not investigate the mechanism for protection, this experiment suggests the potential of ORACLE in the discovery and evolution of a therapeutically relevant function *de novo*.

### Supplementary figures

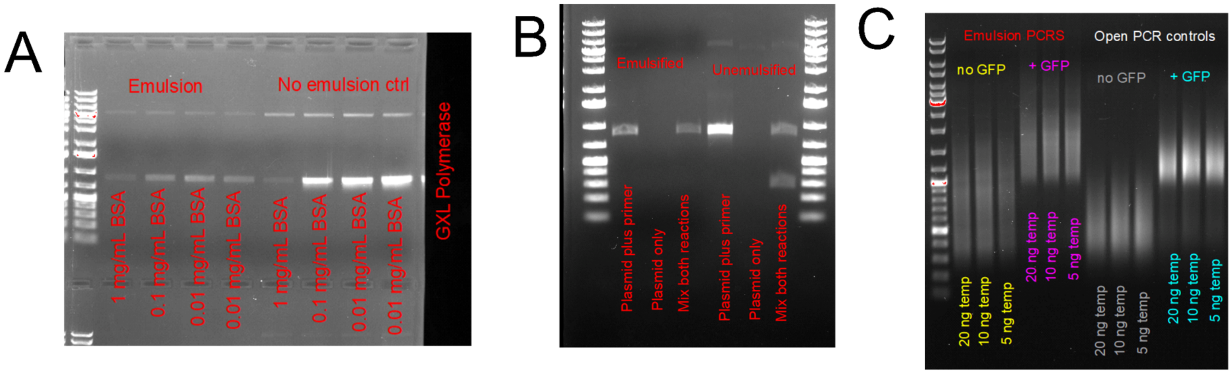


**Fig. S1. Optimization of emulsion PCR and demonstration of decreased size bias in $PCR_{lib}$. (A)** Optimization of BSA concentration in emulsion PCR vs. open control for Takara GXL Polymerase, demonstrating BSA concentrations around 0.01 ng/mL are best tolerated. **(B)** Validation of emulsion stability, two PCRs are prepared with two separate plasmid templates but the same primer binding sites. One has a 700 bp amplicon, one has a 250 bp amplicon. Only the 700 bp reaction contains primers, primers are left out of the 250 bp emulsion. The two separately prepared emulsions are mixed, alongside an open control. The open control demonstrates how both bands are amplified, but the emulsion PCR only yields the larger band indicating the emulsion is stable. **(C)** Representative example of emulsion PCR carried out for the *E. coli* library vs. the open control. Emulsion PCR clearly leads to larger fragments being better preserved.

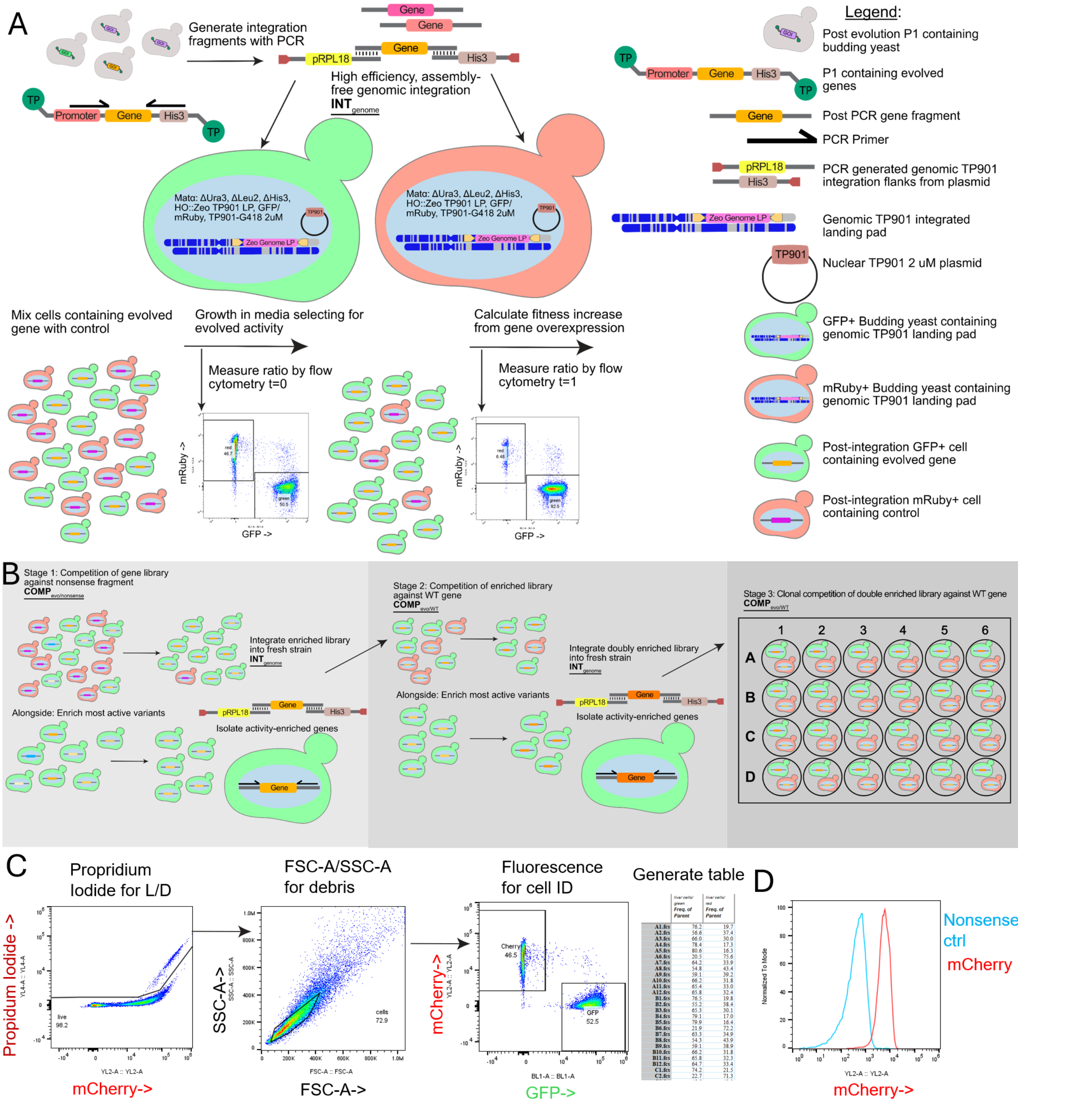


**Fig. S2. Diagram of the transformation method assessment and competition assay setup**. **(A)** Post evolution campaign, DNA samples of interest are extracted with the GC prep method, and the genes are amplified using library specific primers which add overhangs to two genomic integration fragments. Unless otherwise specified the promoter used for all results is pRPL18b. The 5’ fragment contains a TP901 AttB site and the promoter, plus overhangs. The 3’ fragment contains a terminator, *HIS3* marker, and a TP901 AttB site. The overhangs are 60 bp and 55 bp on the 5’ and 3’ respectively. Samples of interest are transformed into strains containing a genomically integrated zeocin AttP TP901 landing pad, and nuclear TP901 overexpression 2µ plasmid, differing only by the integration of an mRuby overexpression cassette, or GFP overexpression cassette (**INT**$_{genome}$). The two fragments are pooled, and co-transformed with the amplified gene fragment, no assembly is necessary, and the integration should yield approximately 100,000 CFU for 50 µL of frozen comp cells. After outgrowth, strain expressing the gene of interest and nonsense control are mixed and taken to competition in selective media to assess impact of evolved genes. **(B)** The assessment proceeds in 3 stages. First, the evolution well is used as a competitor against a nonsense fragment to assess whether there is any activity in evolution well ($\mathbf{COMP}_{evo/nonsense}$), and alongside, the library is passaged in the same selective condition to enrich. If the competition demonstrates activity, the unmixed, enriched library is used as a PCR template to amplify the evolved genes from the integration locus. At this stage, if the evolution well contained multiple active genes, they are gel extracted and sequenced to determine their identity. In stage two, the WT version of the gene is included as a competitor to the evolved sequence ($\mathbf{COMP}_{evo/WT}$). The source of the WT gene varies based on the library (see **Supplementary Text 3**). The enriched gene library, and WT gene are again transformed into competition strains, and the enrichment is repeated. The competition is generally carried out for evolved gene vs. WT, and both evolved vs. nonsense and WT vs. nonsense. If at stage two the evolved variants outcompete the WT gene, the now twice enriched library is again PCR amplified, and retransformed for clonal competition, which is streaked to single colonies. Single colonies are then assessed against the WT gene ($\mathbf{COMP}_{evo/WT}$) and sequenced again. For the most interesting evolution results the clonal sequences were used for further characterization in reporter assays. **(C)** Sample gating and analysis for competition experiments. Wells are first gated for propidium iodide (PI) negativity, keeping in mind that mRuby overlaps with PI signal. They are then gated for size, and then fluorescent populations are counted and exported to tables. **(D)** Demonstration of purity of genomic integration. In place of an evolved gene an mCherry cassette with identical overhangs to the yeast gene library was amplified with the same primers and transformed in the unassembled method, alongside a nonsense control. Only one mode is evident in the mCherry three-part transformation, indicating a high purity of correct integration events, rather than a detectable fraction receiving only the *HIS3* fragment.

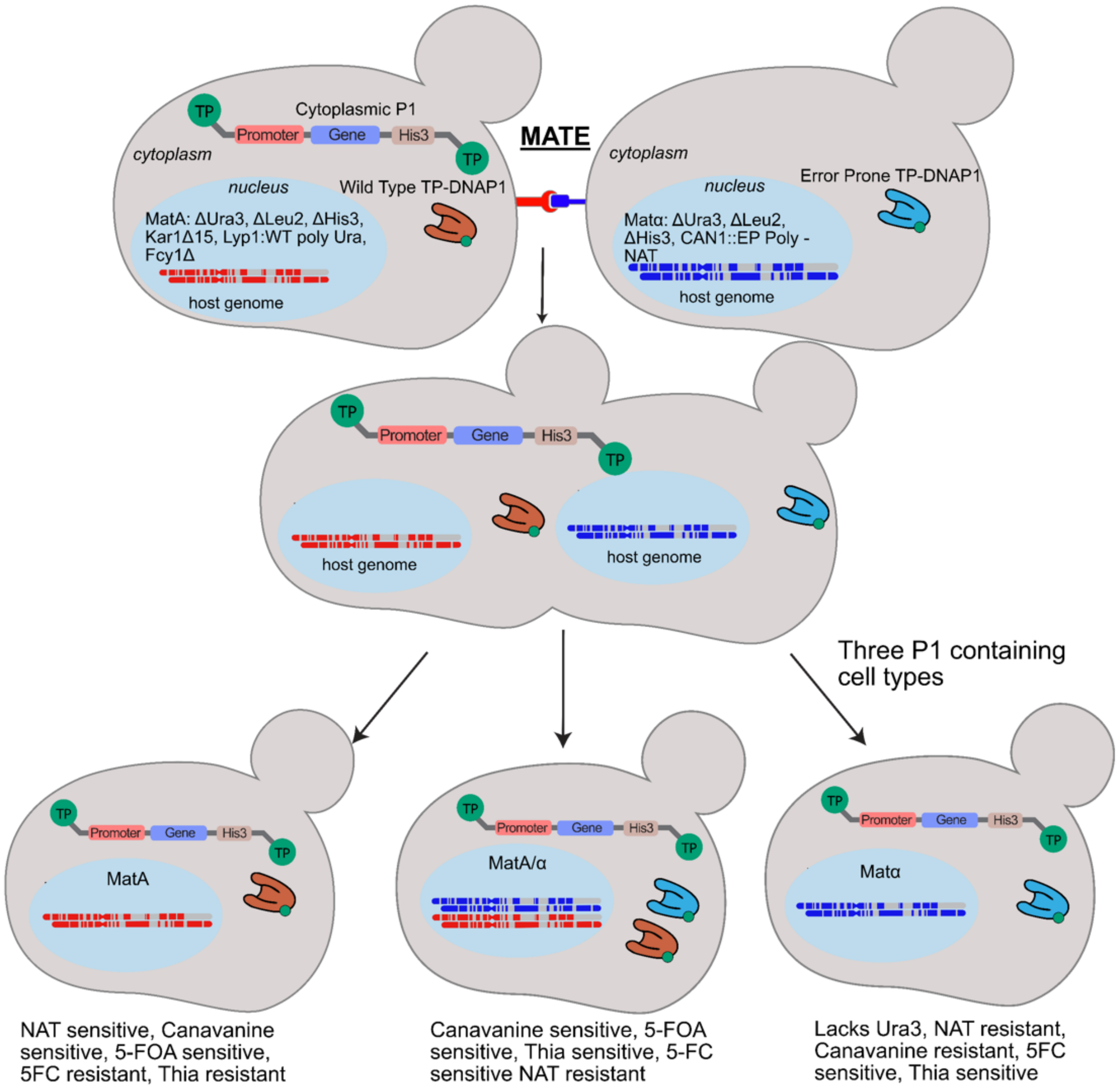


**Fig. S3. Visual representation of the abortive mating technique MATE.** A p1 donor strain mates with a p1 receiver strain, creating 3 cell types with selectable sensitivities. In this example, selection for the p1 receiver strain (*MATα*) would be carried out by selecting for histidine prototrophy, nourseothricin resistance, canavanine resistance, 5-FOA resistance. If mating were in the opposite direction, we would select for the *MATa* strain by using histidine prototrophy, uracil prototrophy, thialysine resistance, and 5FC resistance. This technique with two levels of counterselection reduces the diploid frequency to <1/10,000, which may be sufficient for many selections.

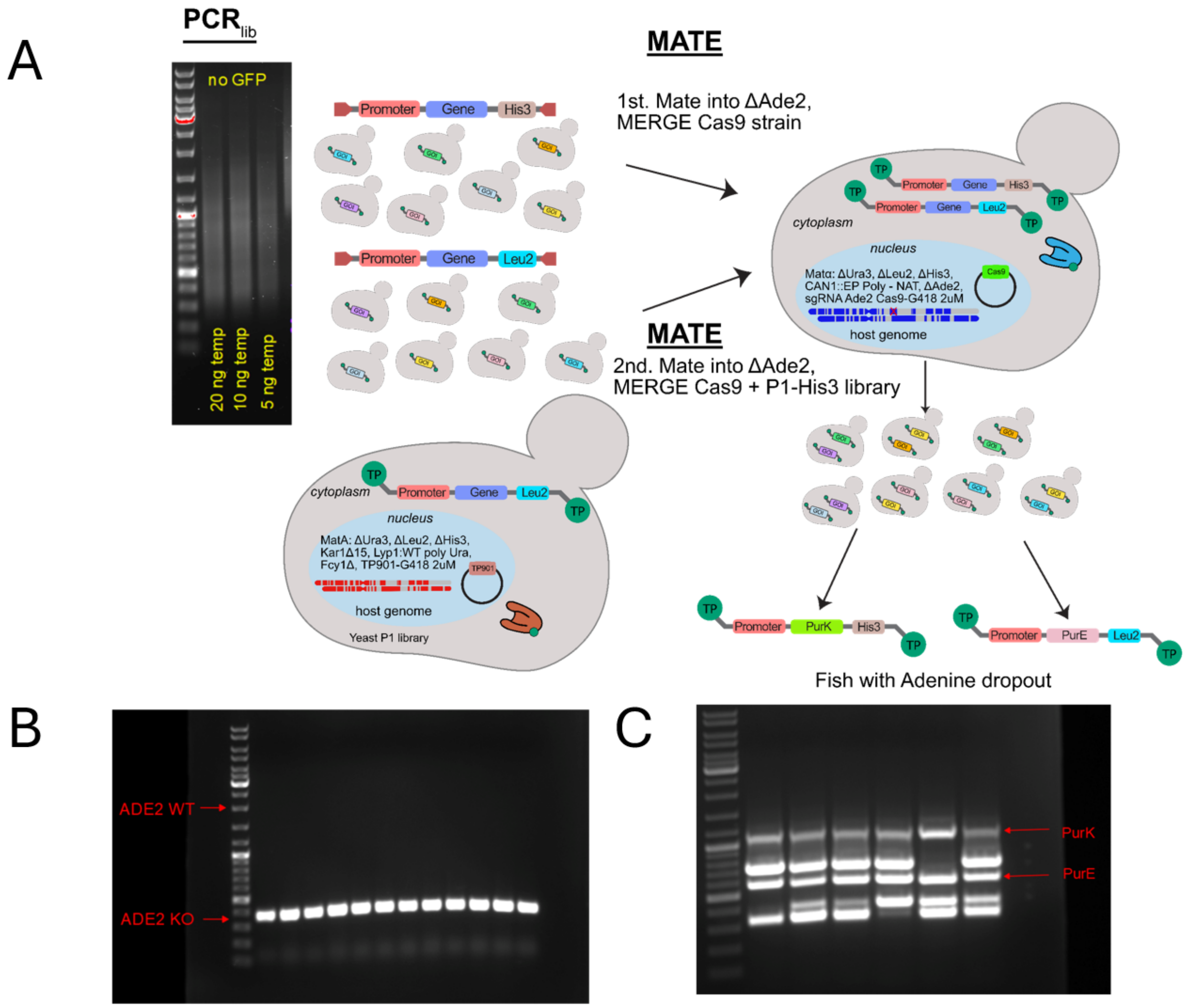


**Fig. S4. Demonstration of ability to fish out a specific combination of genes at a frequency of <1/10,000,000. (A)** Schematic, *E. coli* genes are cloned into two separate integration cassettes, marked with *HIS3* and *LEU2*, and used to generate two p1 donor libraries. These are sequentially mated into a strain lacking genomic *ADE2* and containing a Cas9 plasmid which targets *ADE2*. The use of a Cas9 plasmid to convert the loci of a diploid resulting from mating has been previously described as MERGE (*95*). After counterselection against diploids and selection for the Cas9 plasmid, the cultures were plated, selecting for adenine prototrophy. **(B)** 12x adenine prototrophic colonies were assessed for their genomic copy of *ADE2*, none contained genomic *ADE2*, indicating the rate of diploids forming was less than the detectable limit. **(C)** p1 encoded gene of six colonies was assessed, and the correct two *E. coli* genes are identified in all. Note that the colonies also contained other random p1s. This is explained by incomplete segregation of multiple per cell integrations. No cheaters were detected, demonstrating efficiency of MERGE in depleting genomic copies of diploids which survive counter selection.

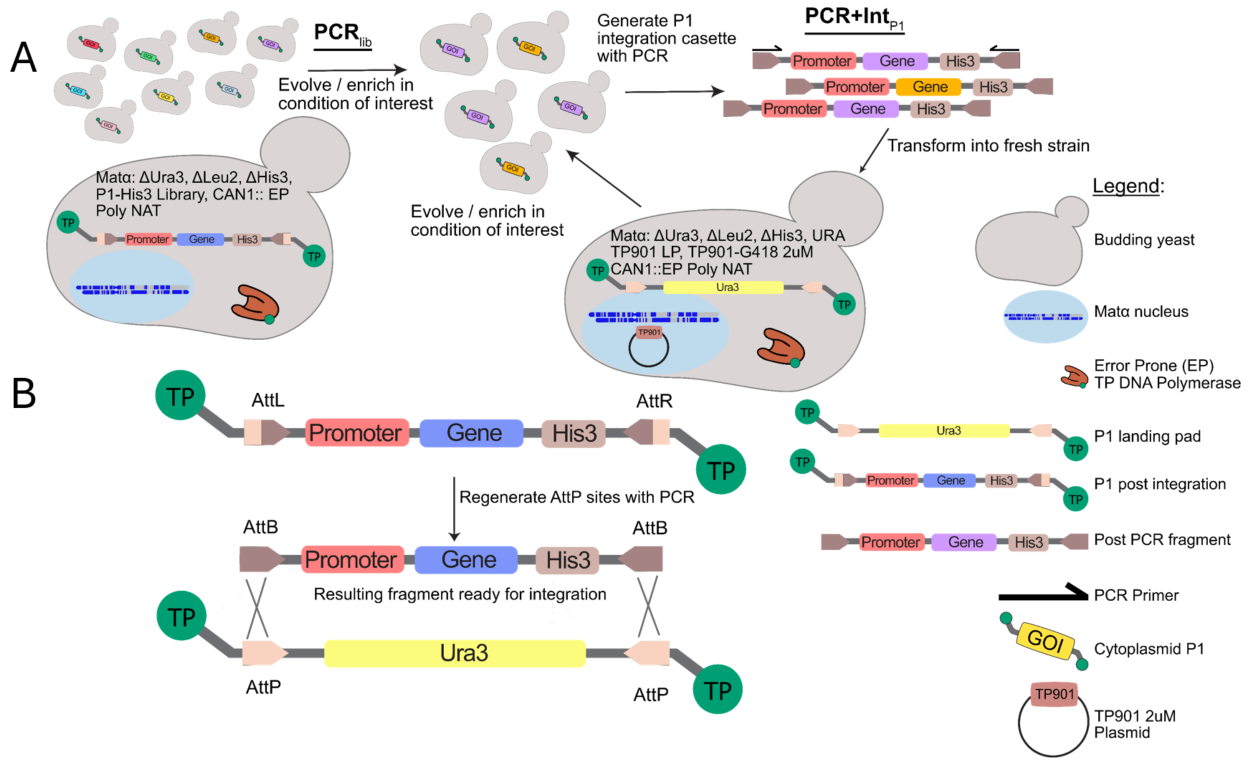


**Fig. S5. Schematic of PCR$_{lib}$ + INT$_{p1}$. (A)** AttB flanked gene library is transformed directly into an error-prone polymerase containing cell with AttP p1 landing pad producing AttL/R flanked cassettes as previously described (*80*). Yeast containing the p1 encoded gene library are passaged several times to enrich / evolve genes for condition of interest. At desired timepoints, DNA is isolated from yeast with GC prep and PCR amplified with primers that regenerate AttL/R into AttB. The resulting library is transformed into fresh error-prone polymerase containing cells. **(B)** Schematic of conversion of AttL/R p1 to AttB for retransformation.

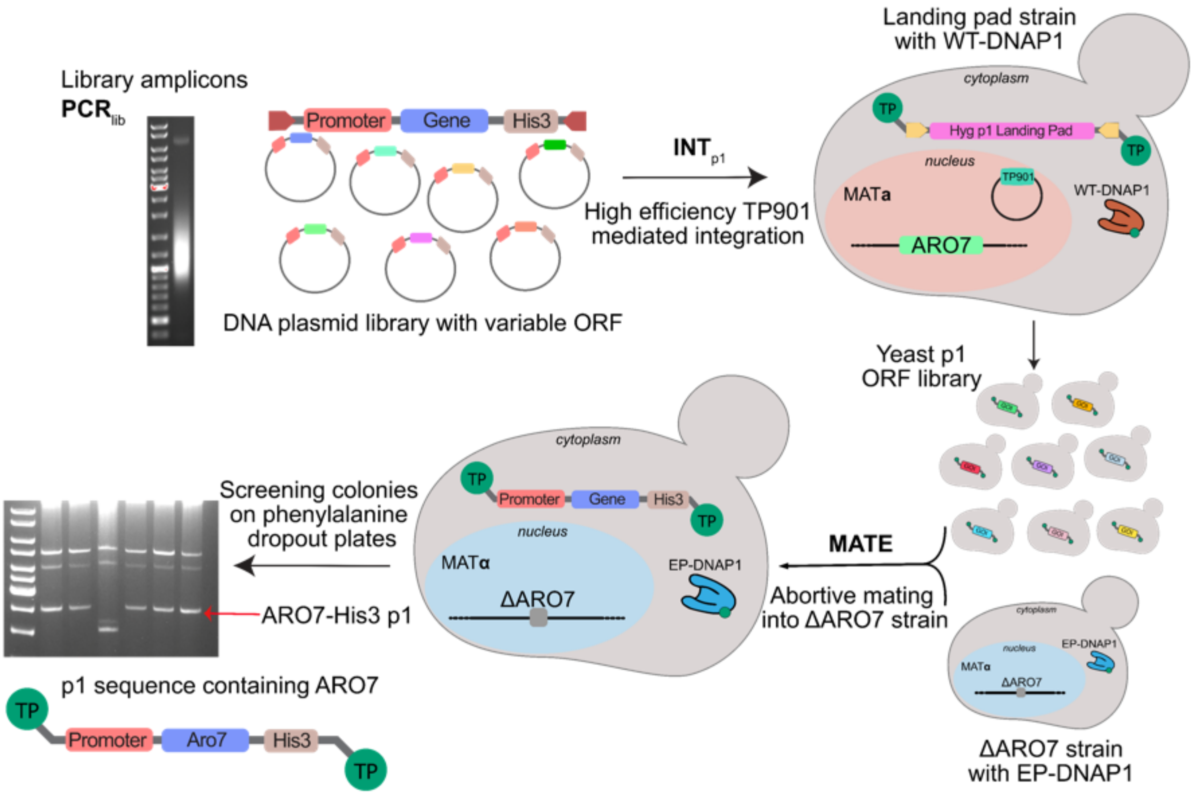


**Fig. S6. Control experiment for the isolation of a positive control gene in ORACLE.** A plasmid library of yeast ORFs (*41*) was subjected to **PCR**$_{lib}$ followed by **INT**$_{p1}$ into a strain that expresses a WT OrthoRep DNAP for p1 replication. Using **MATE**, the p1-encoded library was then transferred into an *Δaro7* selection strain that encoded an error-prone OrthoRep DNAP, resulting in a yeast ORF library where the ORFs continuously hypermutate to support **EVO**. To test whether *ARO7* from the p1-encoded hypermutating yeast ORF library can be isolated by selection, we plated the **MATE** produced library on media lacking phenylalanine (-F) and screened colonies by p1 miniprep and sequencing (see **Supplementary Text 2**). Over the course of two experiments, 9/10 colonies screened for their p1 encoded gene contained the expected *ARO7*, indicating that the **MATE** regime produced >99.99% correct mating events, as any diploid or incorrect haploid (**Fig. S3**) would have contained the genomic copy of *ARO7* from the donor cell. Shown is an agarose gel analysis of six minipreps (*39*) of DNA from yeast able to grow on -F. 5/6 contain the expected p1-*ARO7* plasmid.

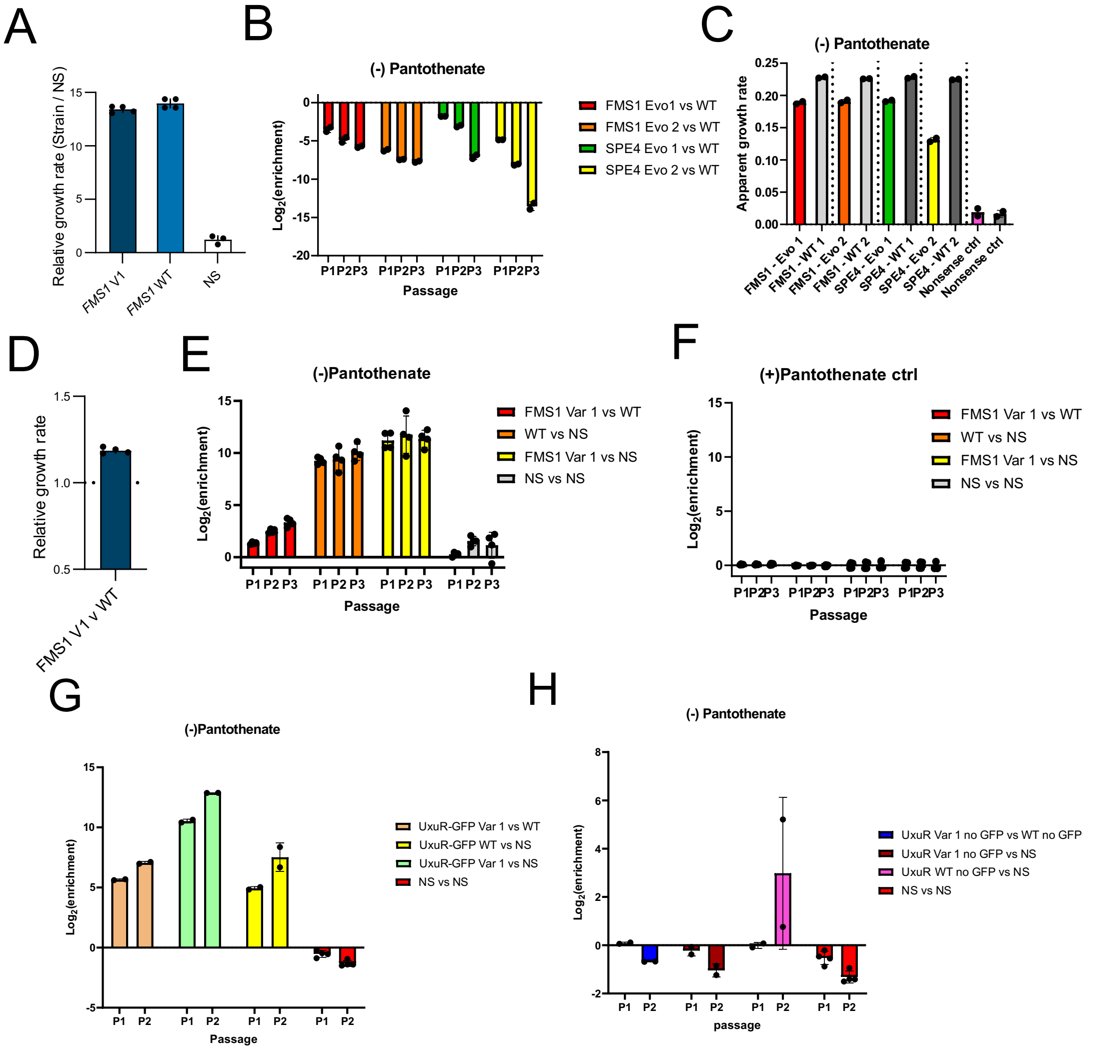


**Fig. S7. Full competition data from the pantothenate evolutions (*Ev-Tol-Pan*). (A-C)** Competition assay data in pantothenate dropout media for high activity pantothenate synthesis enzymes *FMS1* and *SPE4* under the control of pRPL18B promoter. While all evolved and WT enzymes outperform nonsense control, the evolved variants do not confer a growth advantage over their WT counterparts at this expression level. **(D-F)** Competition of the evolved *FMS1 Variant 1* containing mutations V10A, K428R, I481V, A88V against the WT *FMS1* and the nonsense control under the control of the weaker pREV1 promoter in media lacking pantothenate demonstrating the enhanced fitness conferred by the evolved variant over WT (D, E). In media containing pantothenate, there is no fitness impact (F). **(G)** Competition data from the evolved *E. coli UxuR-GFP* Variant 1 against the WT *UxuR-GFP* and both against nonsense controls in media lacking pantothenate demonstrating improvement of *UxuR* by evolution (**Fig. 2A**). **(H)** Competition data from *UxuR* evolved and WT *UxuR* PCR modified to remove the GFP from the ORF, demonstrating lack of activity without a GFP attached (**Fig. 2A**). $Log_2$(enrichment) describes the enrichment of one species over the other at different passages (P1, P2, etc.), as described in **Materials and Methods**.

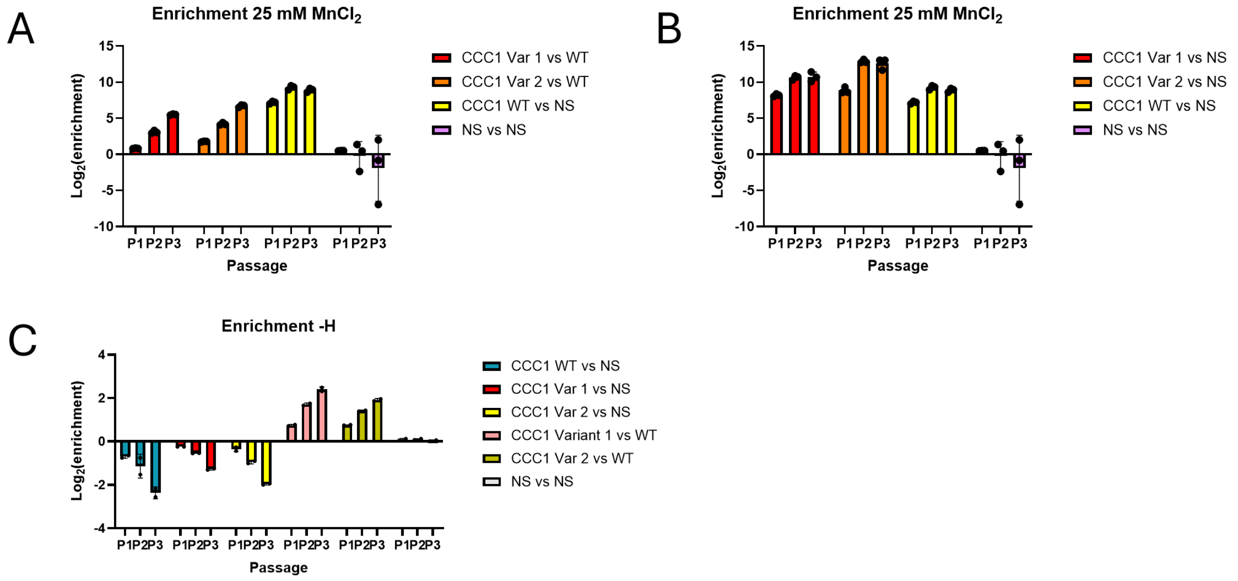


**Fig. S8. Full competition data from the manganese tolerance evolutions (*Ev-Tol-Mn*). (A)** *CCC1 Variant 1* (F92L) and *Variant 2* (F256S) plus (S68P, T274A, A313T) in both, competed against the WT *CCC1*, and WT *CCC1* competed against nonsense demonstrating increase in fitness in toxic (25 mM $MnCl_2$) manganese media, supporting **Fig. 2B**. **(B)** Evolved and WT variants of *CCC1* competed against nonsense control. **(C)** Same variants as in (B) but competed against WT and nonsense control in SC media lacking toxic metal. This demonstrates increase in fitness of evolved variants over WT, but a fitness cost associated with overexpression relative to nonsense control, supporting **Fig. 2B**. $Log_2$(enrichment) describes the enrichment of one species over the other at different passages (P1, P2, etc.), as described in **Materials and Methods**.

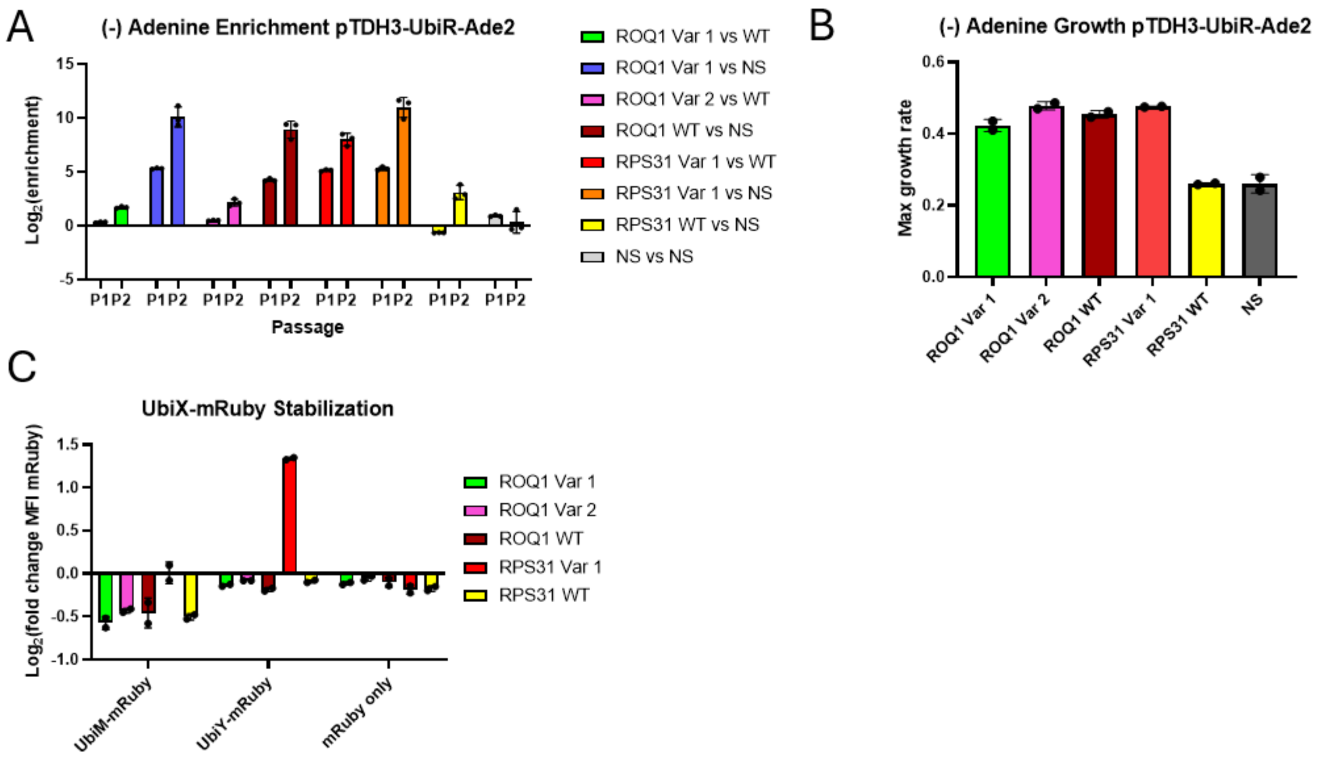


**Fig. S9. Full competition and reporter assay data supporting conclusions made in the main text relating to *ROQ1* and *RPS31* variants (*Ev-Deg-High*). (A)** *ROQ1* variants slightly improved growth rate relative to WT, and *RPS31 R74G* (*Variant 1*) dramatically improved relative to WT *RPS31* which lacks activity (see **Fig. 5A**). $Log_2$(enrichment) describes the enrichment of one species over the other at different passages (P1, P2, etc.), as described in **Materials and Methods**. **(B)** Confirmation of trends from apparent growth rates from competition in a growth curve. Differences in *ROQ1* were too small to reliably detect in a curve, but the *RPS31* difference was confirmed. Reported value is the standard growth rate constant ($hr^{-1}$) as calculated in **Materials and Methods**. **(C)** Fluorescent UbiY-mRuby data demonstrates that *RPS31 Variant 1* increases the level of UbiY-mRuby relative to nonsense, whereas the *ROQ1* variants and WT slightly decreases it. Data reported is the $log_2$ of the relative fluorescence ratio of the sample of interest to that of a nonsense control, measured in biological duplicate transformations.

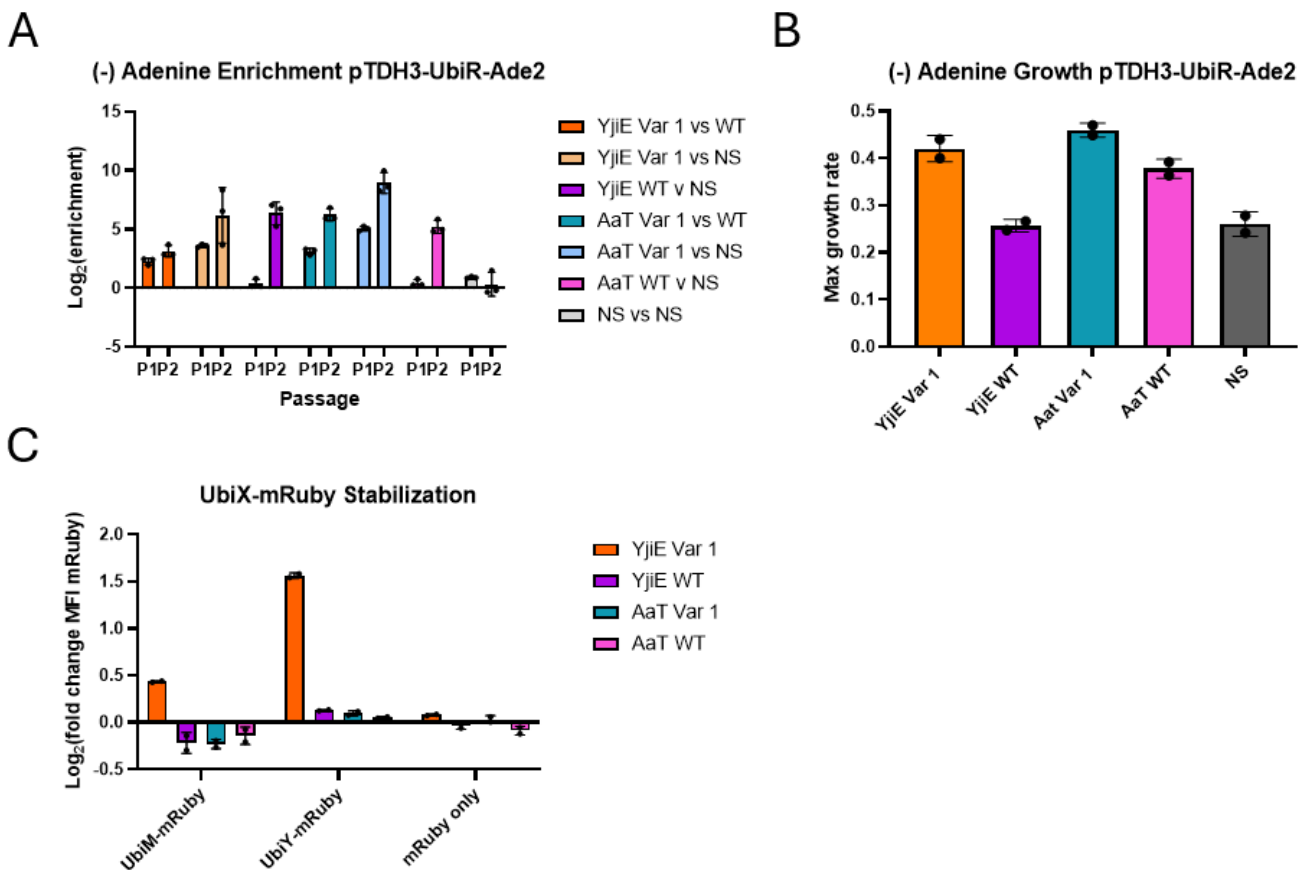


**Fig. S10. Full competition and reporter assay data supporting conclusions made in the main text relating to *YjiE* and *AaT* variants (*Ev-Deg-High*). (A)** *YjiE* F143S (*Variant 1*) significantly improved growth relative to WT which lacks activity, and *AAT* L19P I62V (*Variant 1*) dramatically improved relative to WT AAT which has some basal activity, supporting **Fig. 5A**. $Log_2$(enrichment) describes the enrichment of one species over the other at different passages (P1, P2, etc.), as described in **Materials and Methods**. **B)** Confirmation of trends from apparent growth rates from competition in a growth curve supporting that WT *YjiE* is inactive, WT *AaT* is active, and both evolved variants improved. Reported value is the standard growth rate constant ($hr^{-1}$) as calculated in **Materials and Methods**. **C)** Fluorescent UbiY-mRuby data demonstrating that *YjiE Variant 1* increases the level of UbiY-mRuby, and UbiM-mRuby relative to nonsense, whereas neither *YjiE* WT, nor either measured *AaT* sequences have detectable activity, as expected. Data reported is the $log_2$ of the relative fluorescence ratio of the sample of interest to that of a nonsense control, measured in biological duplicate transformations.

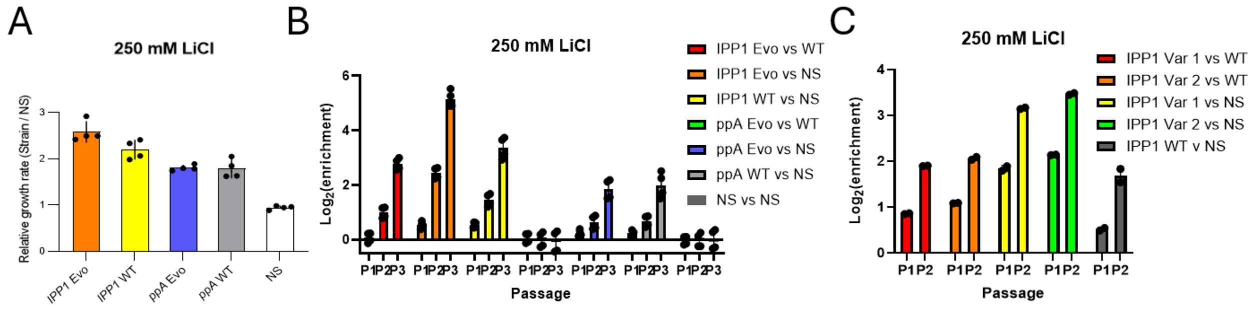


**Fig. S11. Full competition data from the lithium tolerance evolutions (*Ev-Tol-Li*). (A)** Relative growth rates against NS control for bulk yeast *IPP1* variants and *E. coli ppA* variants demonstrating fitness advantages for all variants and WT versions of *IPP1* and *ppA* in media containing a toxic concentration of LiCl (250 mM). Bulk evolved *IPP1* variants conferred additional fitness compared to WT *IPP1*, while *E. coli ppA* variants confer similar fitness compared to WT *ppA*. **(B)** Bulk competition against WT versions of genes and nonsense controls for *IPP1* and *ppA* in 250 mM LiCl media, demonstrating improvement of *IPP1*, and equivalence of *ppA*. Note that the effects of lithium toxicity were apparent mostly after one passage. **(C)** Competition data of individual variants (shared mutations (K11D, I69V, F190L, K199R, T251A,) and variable mutations (*Variant 1*: K228E, *Variant 2*: A232V) against WT and nonsense controls in toxic lithium chloride media, supporting **Fig. 2C**. $Log_2$(enrichment) describes the enrichment of one species over the other at different passages (P1, P2, etc.), as described in **Materials and Methods**.

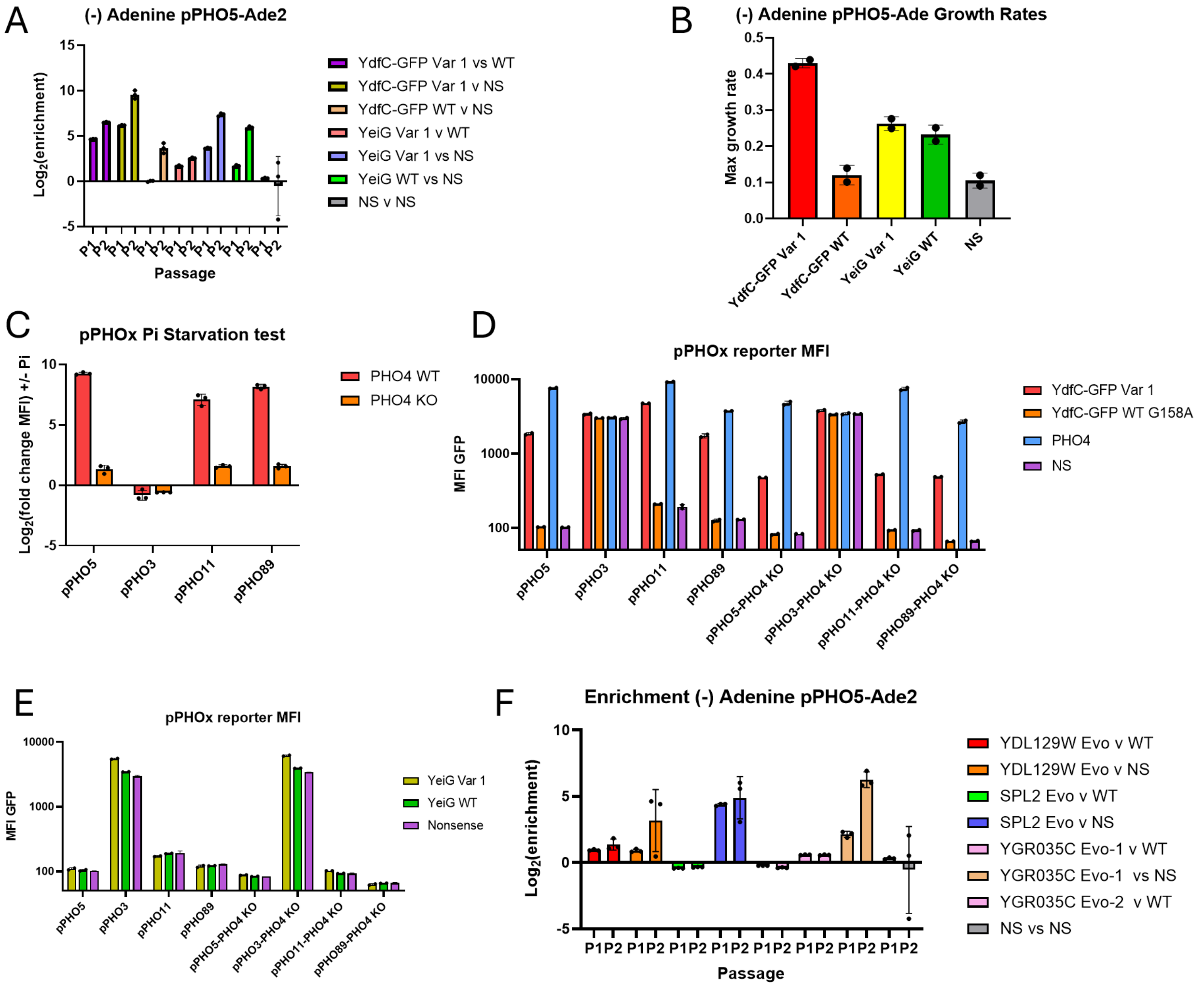


**Fig. S12. Full competition and reporter assay data from the pPHO5 evolutions (*Ev-TF*) supporting Figure 3. (A)** Full competition data in media lacking adenine from the pPHO5-*ADE2* evolutions, notably showing that the *YdfC-GFP* evolved *Variant 1* (D12V, A54V, **GFP** R91H, R165H, H173Y, V204I, R260H, I263T, A298V, V311I, L313H) had activity whereas the WT was largely inactive, and the *YeiG* evolved *Variant 1* (E27G, G89R, A189V, P268S) gained activity over the WT *YeiG*, which did have some activity at the start. This supports **Fig. 3A**. $Log_2$(enrichment) describes the enrichment of one species over the other at different passages (P1, P2, etc.), as described in **Materials and Methods**. **(B)** Remeasured growth rates of noted samples in a standard growth curve, measured in duplicate and calculated maximum growth rate in media lacking adenine, supporting no activity of *YdfC-GFP* WT, high activity with *Variant 1*, and improvement of *YeiG Variant* 1 over WT *YeiG*. Reported value is the standard growth rate constant ($hr^{-1}$) as calculated in **Materials and Methods**. **(C)** Confirmation of reporter assays with no additional genes overexpressed. Cells containing the pPHOx-GFP cassettes, +/- a genomic copy of *PHO4*, were passaged in biotriplicate to phosphate starvation media, and normal SC media for 24 hours, and measured by flow. Data reported is the $log_2$ of relative fluorescence of phosphate starved cells to cells in phosphate containing media ($Log_2$ (fluorescence -$P_i$ / Fluorescence +$P_i$)). **(D)** Non-normalized data of evolved *YdfC-GFP* measured side by side in fluorescent reporter assays against the unevolved sequence and endogenous transcription factor Pho4, demonstrating the activity of the evolved variant and partial maintenance of activity with the endogenous *PHO4* deleted. **(E)** Non-normalized data of evolved *YeiG* measured side by side in fluorescent reporter assays against the unevolved sequence and endogenous transcription factor Pho4, demonstrating the activity of the evolved variant and maintenance of activity with the endogenous *PHO4* deleted. **(F)** Competition data for isolated yeast sequences which contained bioactivity in the pPHO5-*ADE2* context but which failed to improve relative to WT in media lacking adenine. Note that YGR035C was isolated in evolution under both error rates, but did not improve relative to the WT YGR035C. YGR035C and YDL129W are both sequences of unknown functions.

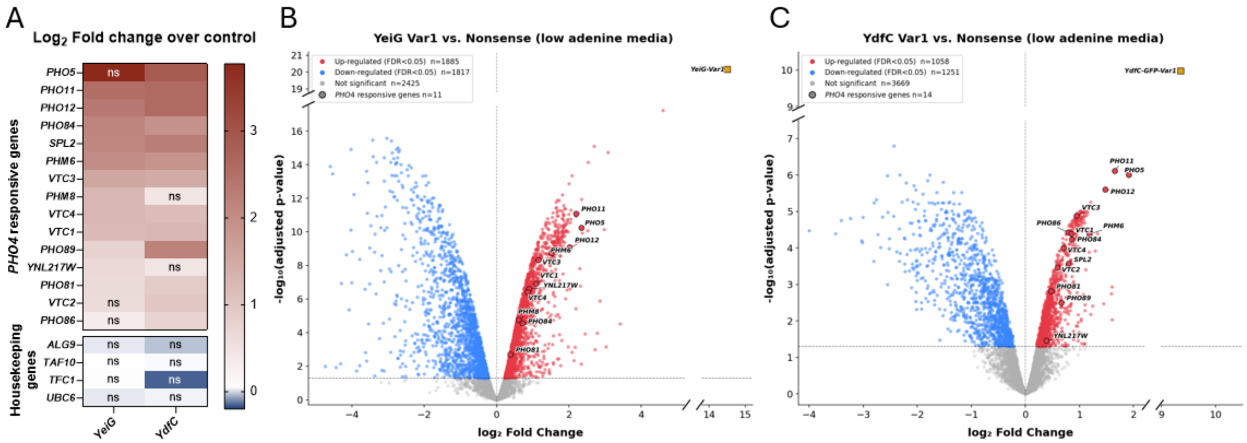


**Fig. S13. The evolved *YeiG* and *YdfC* upregulate the expression of many canonical Pho4-responsive genes. (A)** Heat map showing the gene expression fold change for a set of Pho4-responsive genes and housekeeping genes in excess adenine media (80 mg/L). **(B-C)** Volcano plots showing the RNA-seq data in low adenine media (2.5 mg/L) for (B) *YeiG Variant1*, (C) *YdfC Variant1*, respectively. Volcano plots reveal that Pho4-responsive genes exhibit comparable transcriptional profiles in low-adenine media relative to excess-adenine conditions (**Fig. 3E**). However, due to growth rate differences between the evolved variant and the control strains, expression of thousands of additional genes is affected as well. RNA-seq for *YeiG* and *YdfC* was performed with n = 2 and n =3 biological replicates, respectively. ns, not significant.

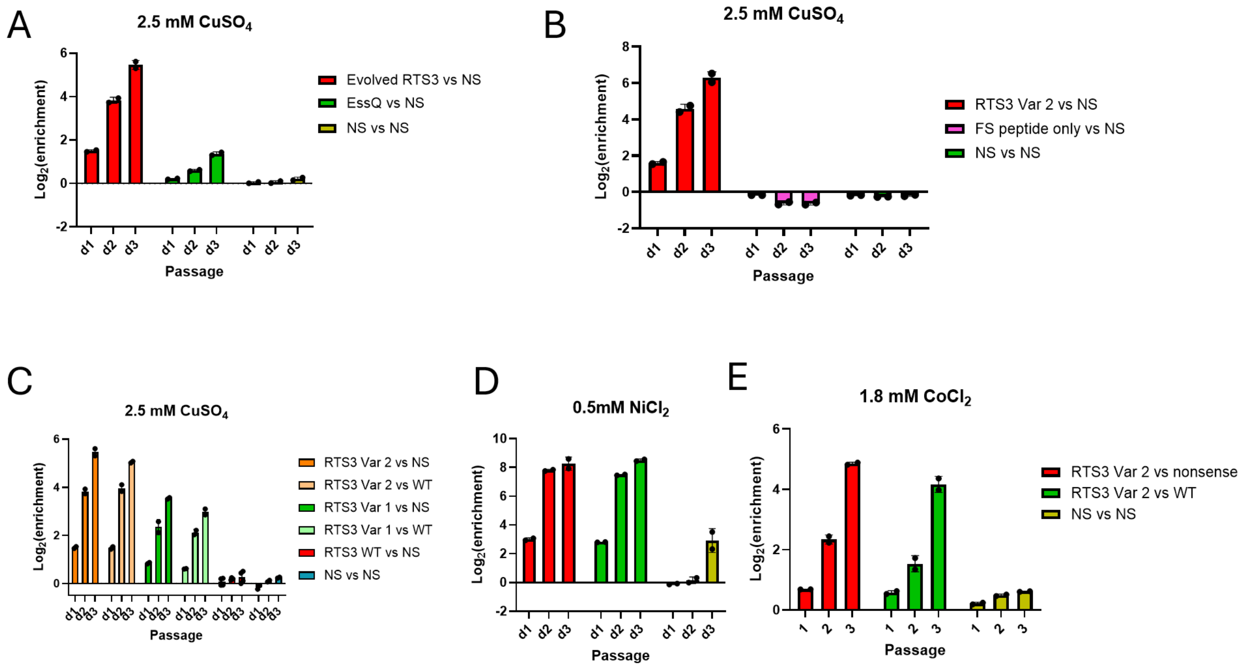


**Fig. S14. Full competition data from the copper tolerance evolutions (*Ev-Tol-Cu*). (A)** Competition data versus nonsense, demonstrating high enrichment of the *RTS3 Variant 2* V8A, A19T, V39I, K74R with a frameshift at residue 102 in 2.5 mM CuSO4, also demonstrating copper resistance by WT *E. coli* EssQ overexpression. *EssQ* is a predicted class II holin of the cryptic *DLP12* prophage (*106*). Its emergence as a gene with activity for copper tolerance is therefore another case of unexpected gene activity. In this case, overexpression of WT *EssQ* increased copper tolerance but the mutant variants produced through **EVO** did not improve fitness of the WT *EssQ* function. **(B)** Competition data of the evolved *Variant 2*, alongside overexpression of the frameshifted peptide alone, demonstrating lack of activity by peptide alone. **(C)** Full competition data supporting the growth rate in **Fig. 2D**. **(D)** Full nickel competition data of *RTS3 Variant 2*, supporting **Fig. 2E**. **(E)** Cobalt competition data supporting **Fig. 2F**. $Log_2$(enrichment) describes the enrichment of one species over the other at different passages (P1, P2, etc.), as described in **Materials and Methods**.

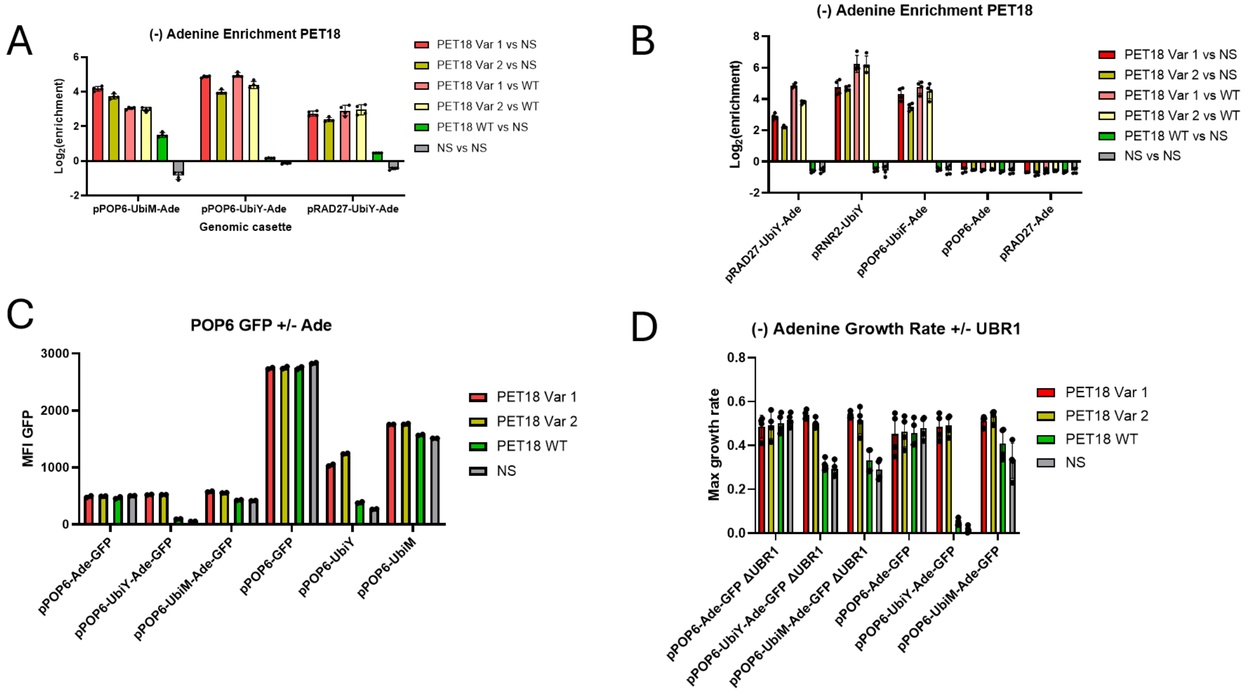


**Fig. S15. Full competition and reporter assay data from the medium strength degron evolutions (*Ev-Deg-Low*) supporting Figure 4. (A)** Single passage competition data in media lacking adenine demonstrating the increased activity of the evolved variants in the context of pPOP6-UbiM/Y, and for another promoter pRAD27-UbiY, demonstrating independence from specific promoter, and activity in UbiM context. $Log_2$(enrichment) describes the enrichment of one species over the other at different passages (P1, P2, etc.), as described in **Materials and Methods**. **(B)** Single passage competition data in media lacking adenine demonstrating activity with another promoter, pRNR2, and with pPOP6-UbiF. Also demonstrating a lack of activity with weak promoters but no degron, demonstrating dependence on presence of degron. $Log_2$(enrichment) describes the enrichment of one species over the other at different passages (P1, P2, etc.), as described in **Materials and Methods**, where each passage represents 24 hours of growth. **(C)** Geomean fluorescence of pPOP6-GFP reporter cells with overexpression of *PET18* variants, demonstrating stabilization of both UbiY, and UbiM, and dependence on the presence of degradation tag for stabilization rather than attachment to Ade2. **(D)** Max growth rates as measured in growth curves in biological and technical duplicate, demonstrating that the increase in growth rate is not entirely dependent on the inhibition of Ubr1 as there is still a growth rate increase for pPOP6-UbiM-*ADE2*-GFP in Δ*ubr1* cells. Reported value is the standard growth rate constant ($hr^{-1}$) as calculated in **Materials and Methods**.

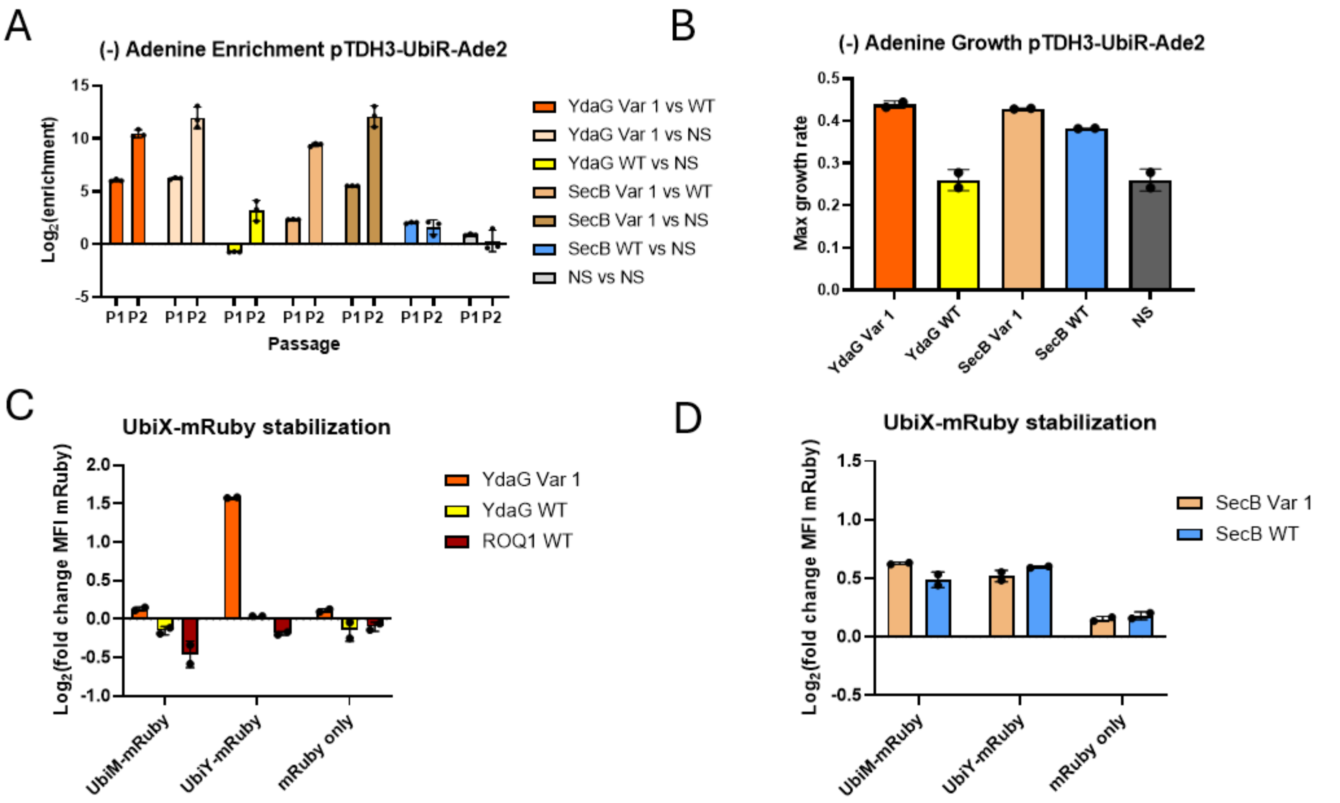


**Fig. S16. Full competition and reporter assay data supporting conclusions made in the main text relating to *YdaG* and *SecB* variants (*Ev-Deg-High*). (A)** *YdaG Variant 1* (A17T, V18M, C30Y, V38M) significantly improved growth relative to *YdaG* WT which lacks activity, and *SecB* with mutations A14T, A41T, Q49L, D59N, V80I, and A145T improved somewhat relative to *SecB* WT which has some basal activity (**Fig. 5A**). $Log_2$(enrichment) describes the enrichment of one species over the other at different passages (P1, P2, etc.), as described in **Materials and Methods**.**(B)** Confirmation of trends from apparent growth rates from competition in a growth curve supporting that WT *YdaG* is inactive, WT *SecB* is active, and both evolved variants improved. **(C)** Fluorescent UbiY-mRuby data demonstrating that *YdaG* A17T, V18M, C30Y, V38M increases the level of mRuby relative to nonsense, whereas the *UBR1* reprogramming *ROQ1* slightly decreases it, and *YdaG* WT is inactive. Data reported is the $log_2$ of relative fluorescence of the sample of interest transformed alongside a nonsense control, measured in biological duplicate transformations. **(D)** Confirmation that *SecB* evolved and WT are not differentiable in the mRuby assay. Data reported is the $log_2$ of relative fluorescence of the sample of interest transformed alongside a nonsense control, measured in biological duplicate transformations.

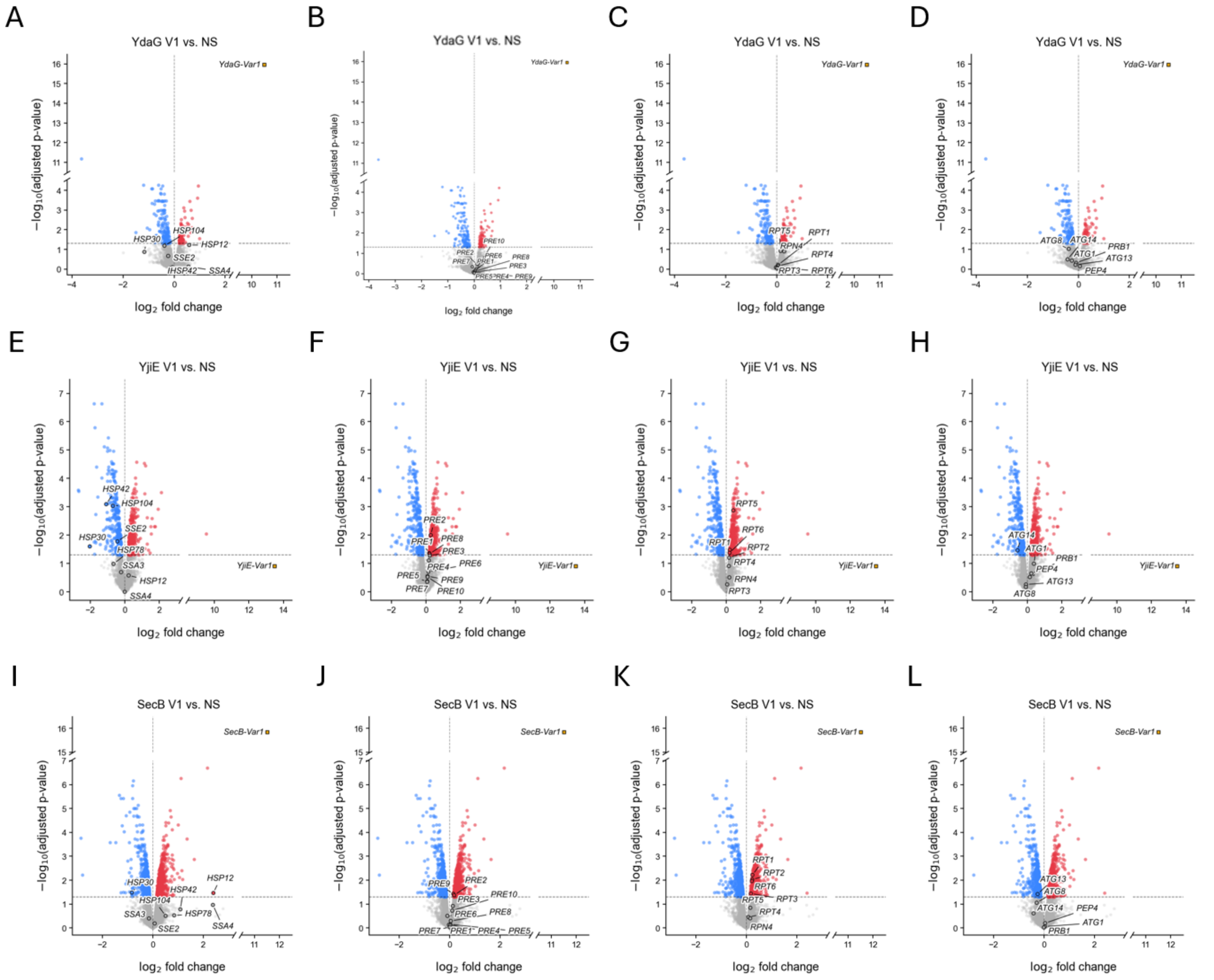


**Fig. S17. RNA-seq data demonstrate the lack of misfolded protein response induction.** Volcano plots representing a list of the most strongly induced heat shock proteins during proteotoxic stress (*74*) (*HSP12*, *HSP104*, *HSP42*, *SSA3*, *SSA4*, *HSP78*, *SSE2*, *HSP30*), proteasome 20S core subunits (*PRE1* - *PRE10*), proteasome 19S regulatory subunits (*RPT1* - *RPT6*), master proteasome regulator (*RPN4*) and some autophagy-related genes (*ATG1*, *ATG8*, *ATG13*, *ATG14*, *PEP4*, *PRB1*) in strains overexpressing *YdaG Variant 1* **(A–D),** *YjiE Variant 1* **(E–H),** or *SecB Variant 1* **(I–L)** compared to nonsense control. The lack of significant upregulation in misfolded protein response genes indicate that these evolved variants are not simply misfolding and overwhelming the degradation machinery. Although a few related genes show minor expression changes in either direction, this sporadic pattern likely reflects the broader impact of *YjiE Variant 1* and *SecB Variant 1* on global protein degradation rather than a coordinated stress response. Gene expression changes were calculated by comparing a strain overexpressing the evolved gene (gold square) to a nonsense control strain. RNA-seq was performed in non-selective adenine-rich (80 mg/L) media with 3 biological replicates.

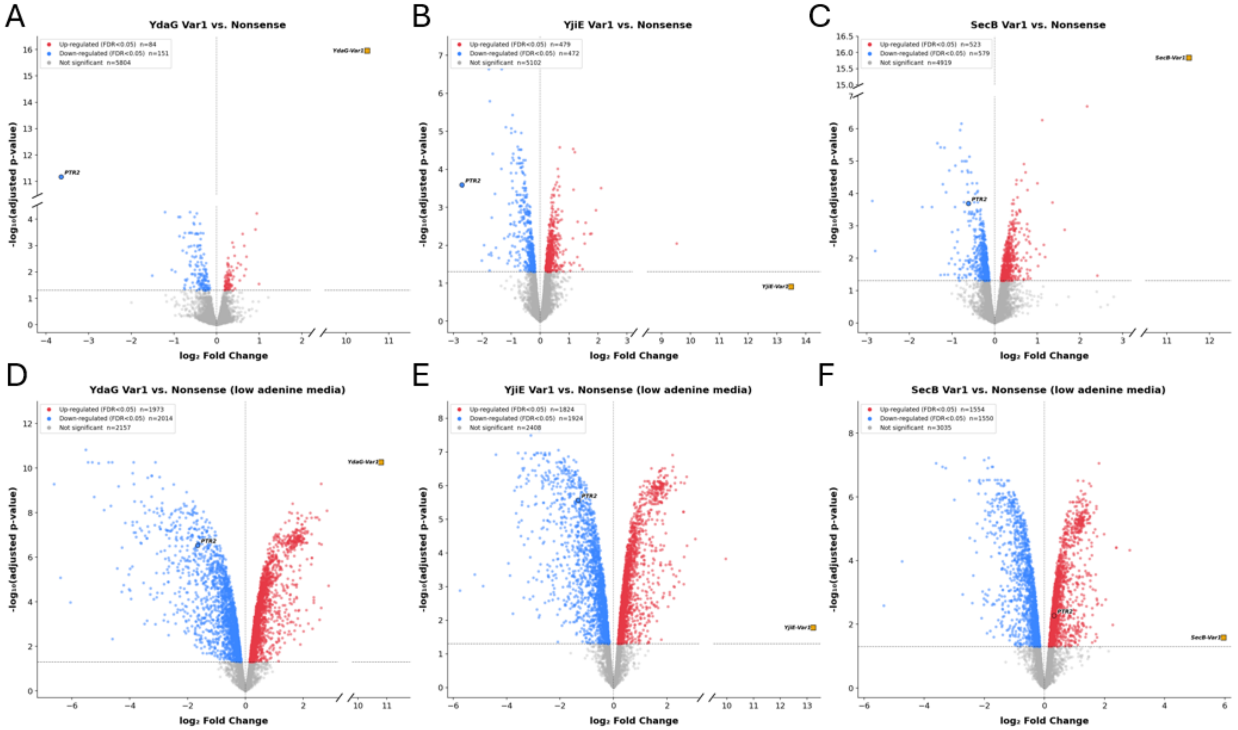


**Fig. S18. RNA-seq data representing *PTR2* expression changes.** Volcano plots illustrating differential gene expression for strains expressing the evolved variants of *YdaG*, *YjiE*, and *SecB* in **(A–C)** adenine-rich (80 mg/L) and **(D–F)** low-adenine (2.5 mg/L) media, respectively. This supports **Fig. 5D**. The *PTR2* gene is shown with a big circle. *YdaG Variant1* and *YjiE Variant1* strongly downregulate *PTR2* while *SecB Variant1* only modestly downregulates (in adenine-rich condition) or upregulates (in low-adenine condition) PTR2. RNA-seq was with 3 biological replicates of each strain.

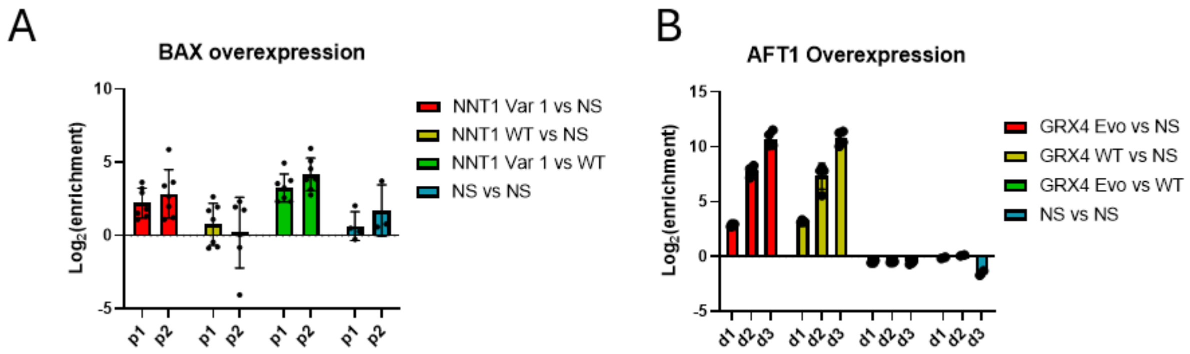


**Fig. S19. Competition data from *BAX* and *AFT1* overexpression selections. (A)** Rescue of *BAX*-induced cell death (see **Supplementary Text 8**). Evolved *NNT1*(D19N, K49R, G60S, G90D, N135D, A219T) significantly enriches over both NS control and WT NNT1 during BAX-induced apoptosis. While *NNT1* WT did not improve growth over the NS control, *NNT1 Variant 1* bearing 6 nonsynonymous mutations improved growth rate significantly. *BAX-2A-URA3* was expressed under an inducible *GAL1 promoter* and growth in uracil-deficient medium selects for the expression of *BAX-2A-URA3.* $Log_2$(enrichment) describes the enrichment of one species over the other at different passages (P1, P2, etc.), as described in **Materials and Methods**. **(B)** Evolution experiment on preventing cytotoxicity caused by the overexpression of *AFT1* (see **Table S1**) yielded *GRX4* from the yeast ORF library. AFT1 is a transcription factor involved in iron homeostasis that has been shown to be highly toxic when expressed at high levels. The wild-type and evolved variants of *GRX4* equally rescued growth in response to *AFT1* overexpression. *GRX4* encodes a glutathione-dependent oxidoreductase that negatively regulates AFT1 promoter binding when coupled with GRX3. Although this interaction between GRX4 and AFT1 is well-studied, it further validates ORACLE's strength in recapitulating known biological interactions.
